# Dual pathway for metabolic engineering of E. coli metabolism to produce the highly valuable hydroxytyrosol

**DOI:** 10.1101/536458

**Authors:** Emmanouil Trantas, Eleni Navakoudis, Theofilos Pavlidis, Theodora Nikou, Maria Halabalaki, Leandros Skaltsounis, Filippos Ververidis

## Abstract

One of the most abundant phenolic compounds traced in olive tissues is Hydroxytyrosol (HT), a molecule that has been attributed with a pile of beneficial effects, well documented by many epidemiological studies and thus adding value to products containing it. Strong antioxidant capacity and protection from cancer are only some of its exceptional features making it ideal as a potential supplement or preservative to be employed in the nutraceutical, agrochemical, cosmeceutical, and food industry. The HT biosynthetic pathway in plants (e.g. olive fruit tissues) is not well apprehended yet. In this contribution we employed a metabolic engineering strategy by constructing a dual pathway introduced in *Escherichia coli* and proofing its significant functionality leading it to produce HT. Our primary target was to investigate whether such a metabolic engineering approach could benefit the metabolic flow of tyrosine introduced to the conceived dual pathway, leading to the maximalization of the HT productivity. Various gene combinations derived from plants or bacteria were used to form a newly-inspired, artificial biosynthetic dual pathway managing to redirect the carbon flow towards the production of HT directly from glucose. Various biosynthetic bottlenecks faced due to *feaB* gene function, resolved through the overexpression of a functional aldehyde reductase. Currently, we have achieved equimolar concentration of HT to tyrosine as precursor when overproduced straight from glucose, reaching the level of 1.76 mM (270.8 mg/L) analyzed by LC-HRMS. This work realizes the existing bottlenecks of the metabolic engineering process that was dependent on the utilized host strain, growth medium as well as to other factors studied in this work.

## Introduction

Hydroxytyrosol (3,4-Dihydroxyphenylethanol; HT) is a mono-phenolic compound traced in olive fruits [1] and tissues [2], in extracted olive oil [3], or even in olive mills waste waters [4]. It shows a broad spectrum of biological properties due to its strong antioxidant and radical-scavenging capacity [5]. Contributing most of its qualitative characteristics to olive oil, HT was recently approved as a part of “olive oil healthy polyphenols” by EC Regulation 432/2012 [6]. Concisely, the most important properties that make HT so attractive are a) its ability to scavenge the free radicals and thus acting as a potent antioxidant [7], b) the ability to reduce the risk of coronary heart disease and atherosclerosis [8, 9], c) the ability to prevent the LDL oxidation, platelet aggregation, and inhibition of 5- and 12- lipoxygenases [10], d) its critical effects on the formation and maintenance of bones, being used as an effective remedy in the treatment of osteoporosis symptoms [11] and e) its ability to control human and plant bacterial and fungal pathogens [3, 12-15].

In nature, the biosynthetic pathway for the production of HT (e.g. olives or grapes) has not been justified yet [16]. The HT content in olive fruit is highly depended on the cultivar, the cultivation techniques, the environmental conditions as well as the extraction process [17]. Owen and colleagues have comprehensively worked with the phenolic content of olives and virgin olive oil, estimating HT at about 14.4 mg/kg in oil [18, 19] and at 5.8 mg/kg at pericarp tissues [20]. In another study the HT contained in olive oil was estimated to range from 1.38 mg/kg to 7.94 mg/kg [21]. It was found that HT could be recovered from olive mill wastewaters, but low yields and complex extraction processes made recovery too expensive [22]. Interestingly, Agalias et al. [4] attempted and succeeded to purify HT from olive mill wastewaters. On the other hand, obtaining high amounts of HT by chemical synthesis is regarded cost inefficient for industrial scale production. Therefore, alternative cost-efficient strategies to produce HT at considerable high yield are desired [23].

The abovementioned properties along with the fact that it is a compound with extremely high commercial value, make its biosynthesis by biotechnological means attractive. Bioconversion of tyrosol to HT has been reported earlier by Espin et al. [24] using a mushroom tyrosinase (TYR) as a biocatalyst and by Orenes-Pinero et al. [25] by the utilization of a phenol hydroxylase gene from *Geobacillus thermoglucosidasius*. The disadvantages of this approach include the high cost and instability of the TYR enzyme. Later, a soil bacterium identified as *Pseudomonas aeruginosa*, was isolated based on its ability to grow on tyrosol as a sole source of carbon and energy [26]. During growth on tyrosol, this strain promoted the formation of HT and trace quantities of hydroxyphenylacetic acid and 3,4-dihydroxyphenyl acetic acid. Although these methods were successful for the production of HT, they were either cost-inefficient because of the required protocols for enzyme production [24] or required the supplementation of relatively expensive intermediates [26]. On the other hand, Satoh et al. [27] managed to produce HT utilizing an *Escherichia coli* system, achieving however very low titers.

Previous research attempts to produce HT directly from glucose resulted in 12.3 mg/L (0.08 mM) from *E. coli* grown in M9 broth supplemented with yeast extract [27]. They reconstituted the HT pathway utilizing a tyrosine hydroxylase, a biopterin regeneration pathway, a L-3,4-dihydroxyphenylalanine (DOPA) decarboxylase (DDC) and a tyramine oxidase expressed into JW1380 *E. coli* hosts. However recently, Chung et al. [28] followed an alternative approach by utilizing a phenyl acetaldehyde synthase to convert tyrosine into 4-hydroxyphenyl acetaldehyde, which was sequentially converted to tyrosol by an alcohol dehydrogenase (alternate name for aldehyde reductase) and finally to HT by the action of and 4-hydroxyphenylacetate 3-hydroxylase (HpaBC). This approach led to the production of HT directly from glucose at a concentration of 208 mg/L.

In this research work, we describe a strategy for the grafting of an engineered pathway into *E. coli* cells to produce HT through a dual invented pathway (Fig 1). We present a dual approach for the production of HT directly from glucose, managing to score one of the highest HT concentrations produced so far. Specifically, we have tailored the so-called HT pathway with tyrosine as a precursor, utilizing the following genes: an aromatic amino acid decarboxylase (*AADC*), an aldehyde reductase (*ALR*) and a tyrosinase (*TYR*) (Table 1). Various efforts were undertaken to optimize this heterologous HT production, tested under variable genetic and growth conditions such as host selection, fermentation conditions, gene dose, protein expression level, and special enzyme properties. We marked each factor’s specific effect on the metabolic flux and on the final HT yield. To achieve this, we engineered regular strains [BL21(DE3) and HMS174(DE3)], expressing genes not only to produce HT, but also for the endogenous tyrosine overproduction, thus leading to highest titers of HT directly from glucose. Such a metabolic effect, permitted the redirection of primary metabolic flux from externally added glucose to a tyrosine pool that proved to be saturating the first enzymatic step (Fig 1) of the engineered pathway from tyrosine to HT. Moreover, the addition of a selected *ALR (ALR-K*, Table 1), overexpressed along with the other abovementioned genes of the HT pathway (Fig 1B), led to additional increase in HT production efficiency.

**Table 1.**
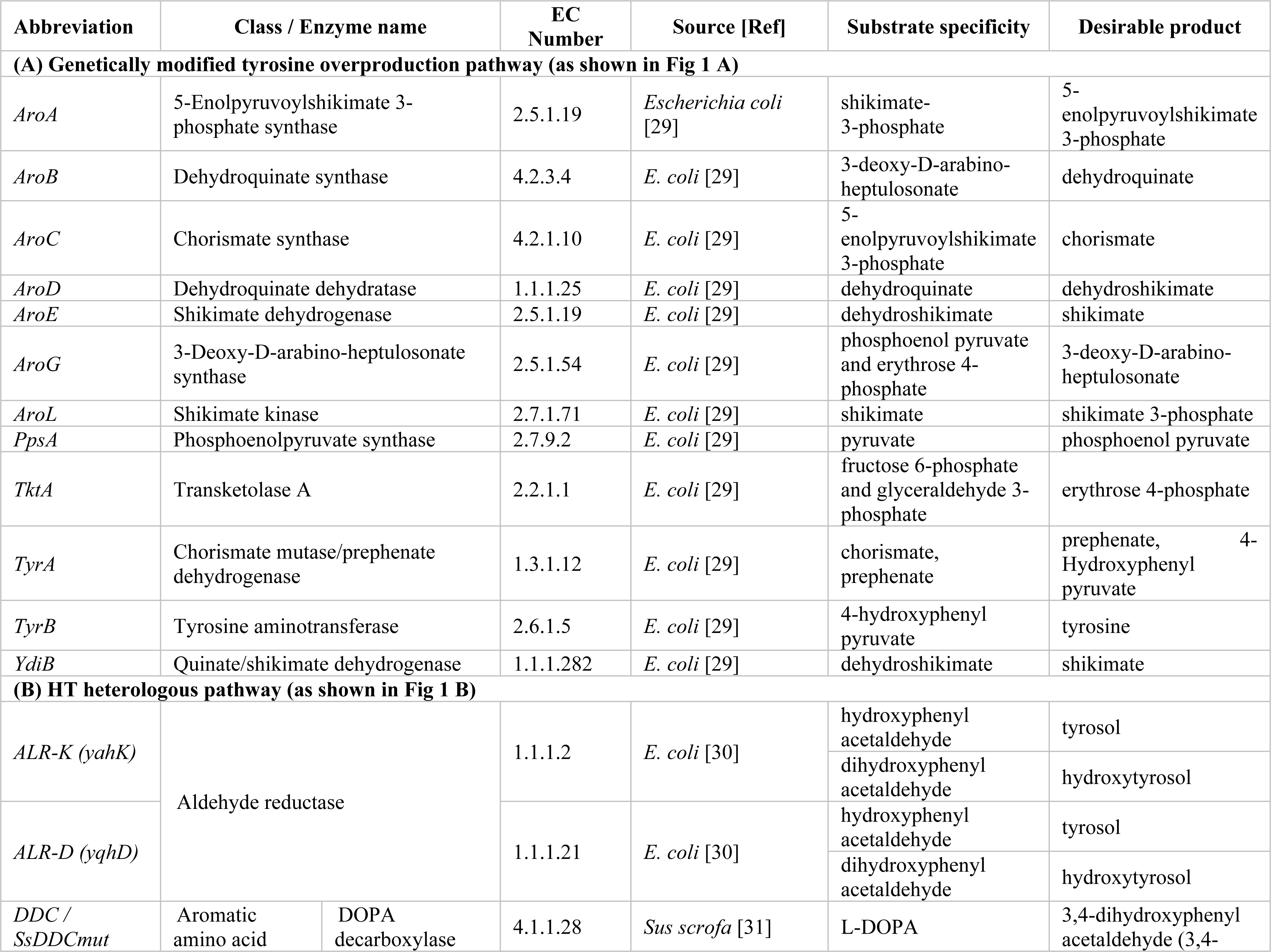
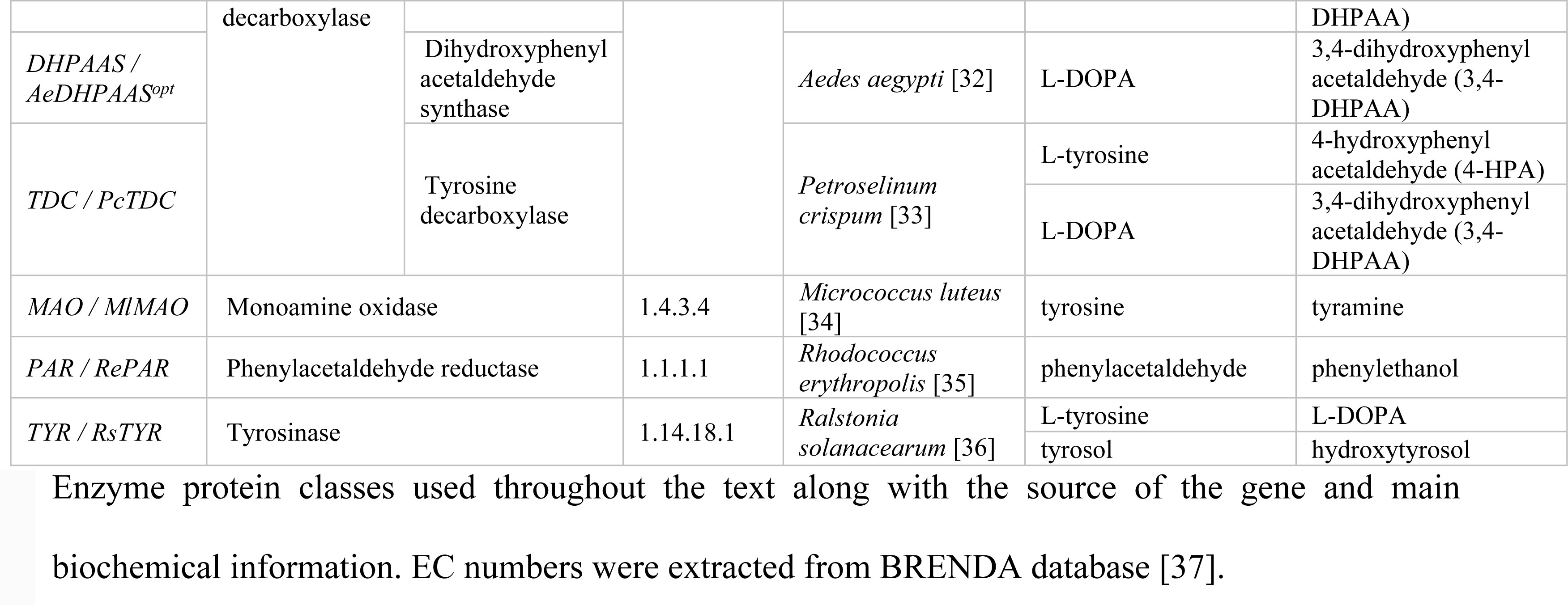
All different gene/enzyme descriptions used in this study.

**Fig 1.**
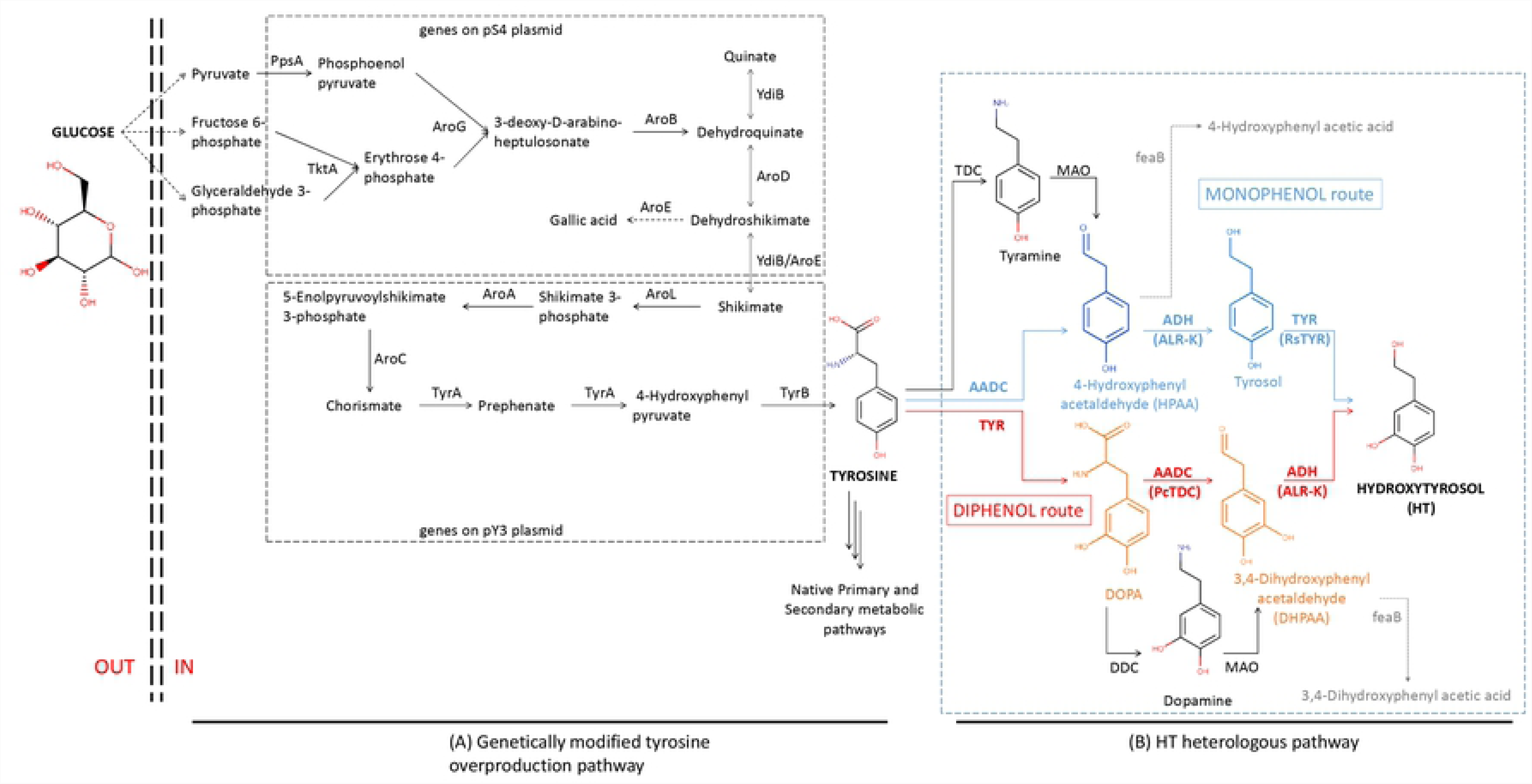
Metabolic scheme for the production of hydroxytyrosol (HT) in *Escherichia coli* directly from glucose. (A) Tyrosine is produced through an overproducing metabolically modified machinery of the primary metabolism (Juminaga et al. 2012) and serves as the main precursor channeled to (B) the newly introduced HT pathway able to transform tyrosine into HT. Double-dashed lines depict *E. coli*’s outer membrane. Dotted arrows depict multiple reaction steps. Due to the dual specificity of the utilized aromatic amino acid decarboxylase (AADC) cloned from parsley, and to tyrosinase (TYR), cloned from *Ralstonia solanacearum*, a dual pathway is generated. The “blue” pathway (monophenol route) portrays the enzymatic route for HT biosynthesis deriving from the decarboxylation of tyrosine, while the “red” pathway (diphenol route) portrays the HT route deriving from the hydroxylation of tyrosine to L-3,4-dihydroxyphenylalanine (DOPA). Gene-respective enzyme abbreviations: *AADC*, Aromatic amino acid decarboxylase. *ALR*, aldehyde reductase. *AroA*, 5-enolpyruvoylshikimate 3-phosphate synthase. *AroB*, dehydroquinate synthase. *AroC*, chorismate synthase. *AroD*, dehydroquinate dehydratase. *AroE*, shikimate dehydrogenase. *AroG*, 3-deoxy-D-arabino-heptulosonate synthase. AroL, shikimate kinase. *DDC*, DOPA decarboxylase. *feaB*, phenylacetaldehyde dehydrogenase, *MAO*, monoamine oxidase. *PpsA*, phosphoenolpyruvate synthase. *TDC*, tyrosine decarboxylase. *TktA*, transketolase A. *TYR*, tyrosinase. *TyrA*, chorismate mutase/prephenate dehydrogenase. *TyrB*, tyrosine aminotransferase. YdiB, quinate/shikimate dehydrogenase (see further details in Table 1). Bold abbreviations denote the genes that were utilized in this study.

## Materials and Methods

### Materials and DNA extraction

All chemicals used were purchased from Sigma-Aldrich unless otherwise stated. Genomic DNA extractions were accomplished from overnight cultures with Qiagen Blood & Tissue kit according to manufacturer instructions. PCR amplified fragments and digested fragments were purified with Macherey-Nagel NucleoSpin Gel and PCR Clean-up kit. Restriction enzymes were purchased from Minotech, Crete, Greece or New England Biolabs. Taq polymerase was provided from Kapa Biotech while the proof-reading polymerase used for all the cloning processes (Phusion) was obtained from New England Biolabs. For SDS PAGE the PiNK prestained protein ladder was used (Nippon Genetics).

### Genetic material, microbial hosts and cloning vectors

All gene descriptions are listed in Table 1. The *TYR* gene was cloned from the strain GMI1000 of *Ralstonia solanacearum* [36], *the AADC gene was cloned from root cDNA of Petroselinum crispum* (parsley). The *yahK* and *yqhD* genes encode for Aldehyde reductases and were cloned from *E. coli* genomic DNA [30]. For further convenience, we will refer to these two genes with the abbreviations *ALR-K* and *ALR-D* respectively (Table 1). DDC gene, encoding for DOPA Decarboxylase from *Sus scrofa*, donated by Prof. C. B. Voltattorni [31]. All genes involved in the tyrosine overproduction pathway (Fig 1A) were kindly donated by Prof. J.D. Keasling [29].

All cloning and sub-cloning steps were done using the Novagen pRSF and pET dual expression (Duet) vectors, all comprising the pET protein expression system. It is developed for the cloning and expression of recombinant proteins in *E. coli*. Target genes were cloned into pET plasmids under the control of a strong bacteriophage T7 transcription and (optionally) translation signals.

The *Escherichia coli* strain used for accomplishing the cloning strategy was the DH10B while the host bacterial strains used for expression experiments were the BL21 and HMS174 (Table 2). BL21 and HMS174 were evaluated for use as hosts because the former belongs to the B line of *E. coli*, while the second belongs to the K12 *E. coli* line [38].

**Table 2:**
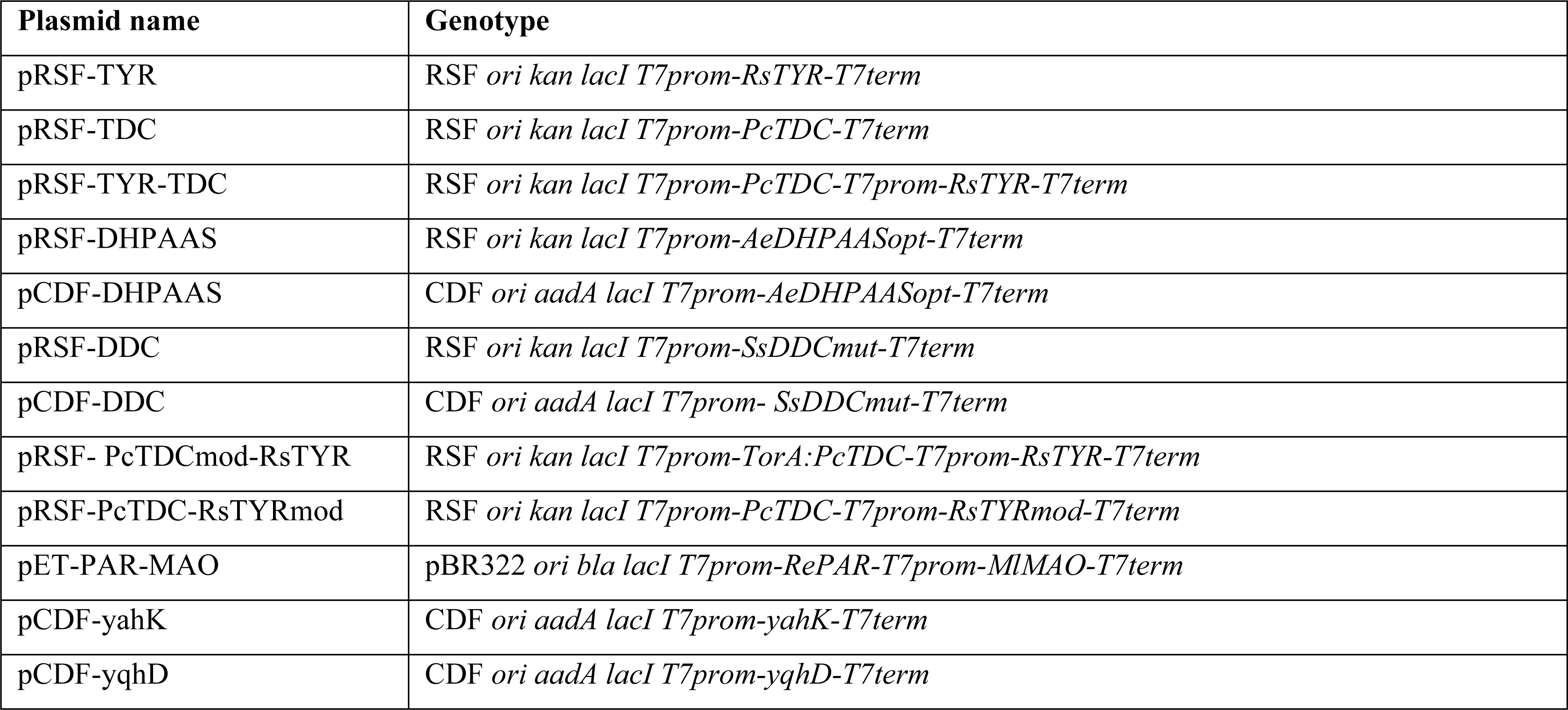
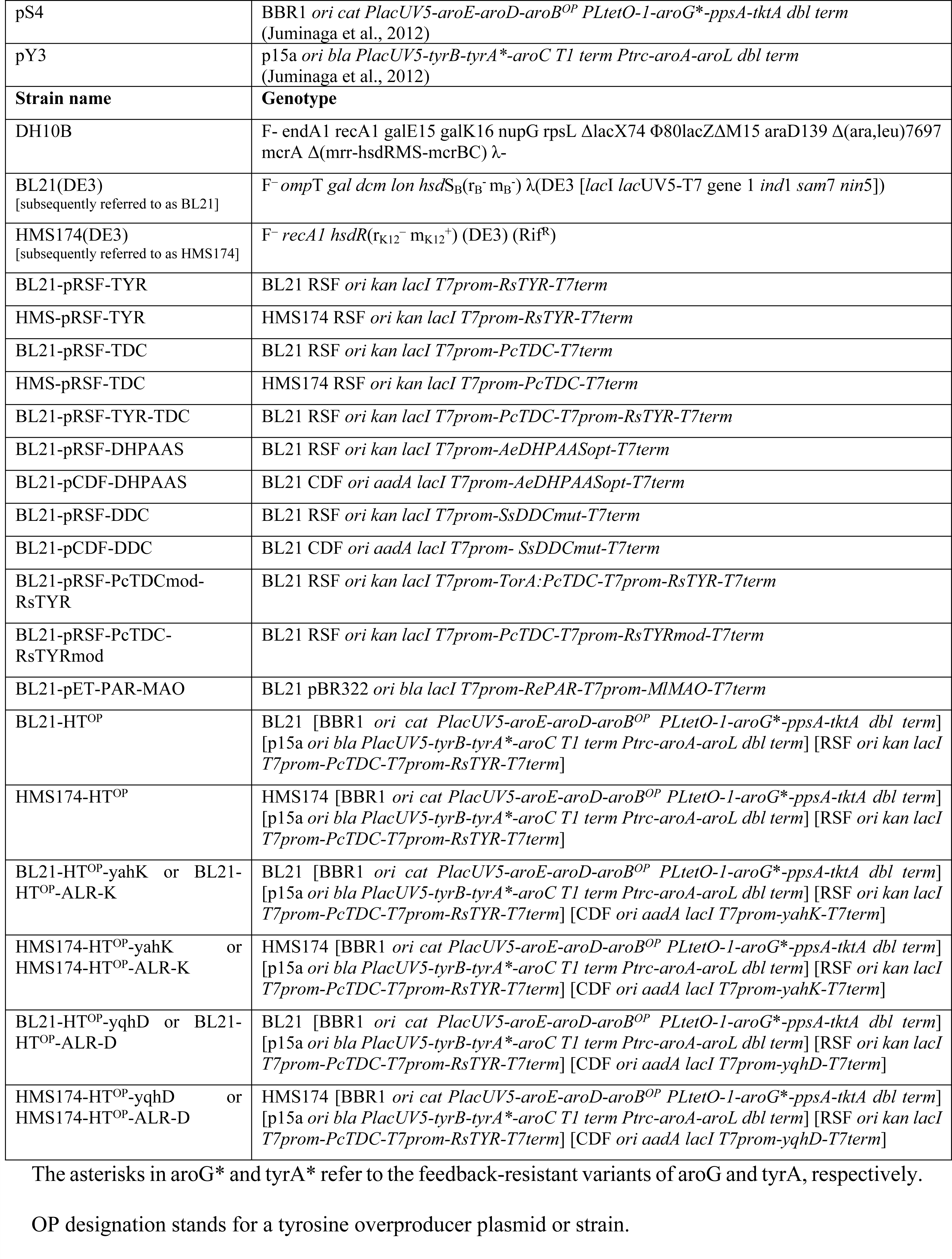
Metabolically engineered bacterial plasmids (see Table 1 for further details) and strains constructed or used in this work.

### Software tools

The prediction of twin-arginine translocation signal peptide in *R. solanacearum* tyrosinase was assessed with the PRED-TAT web integrated tool (ww.compgen.org/tools/PRED-TAT/submit) utilizing Hidden Markov Models [39]. Primer design and *in-silico* DNA manipulations were completed with SnapGene (http://www.snapgene.com/).

### Engineering of expression vectors with genes involved in Hydroxytyrosol biosynthesis

#### Construction of tyrosinase and aromatic amino acid decarboxylase expression vectors

All engineered plasmids constructed in this study were based on Duet expression vectors. The construction strategy and steps for each engineered plasmid used in this study are depicted in Fig 2. Genes were either cloned by PCR utilizing appropriately designed oligonucleotides from various sources (Table S1), or artificially synthesized (Table S1). To confirm each gene’s identity the cloned PCR fragments were sequenced and compared *in silico* with the NCBI deposited sequences.

**Fig 2.**
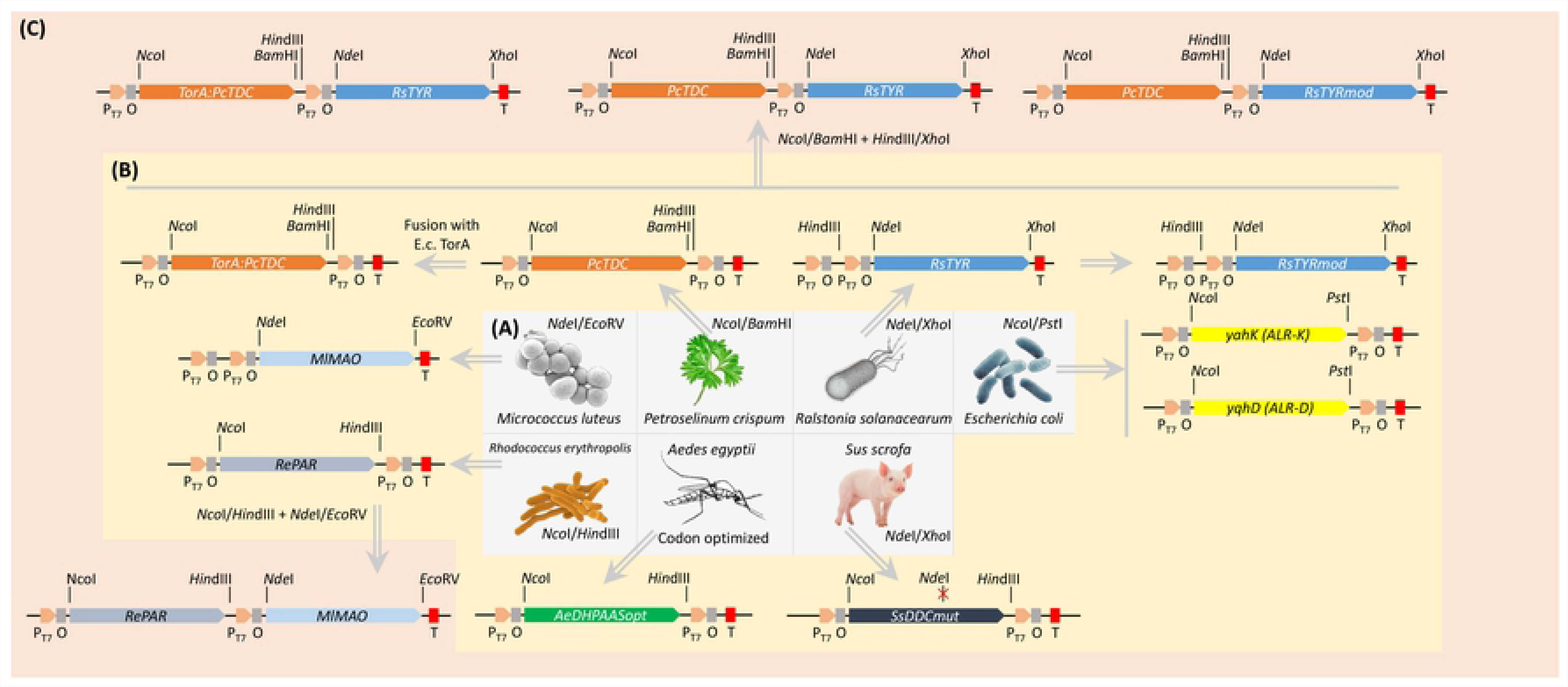
Schematical representation of the cloning strategy to construct the DNA plasmids used throughout this work. Three levels of constructions were created (A), (B), (C). Level (A) deals with the DNA products generated either by PCR using the appropriate primers or by direct synthesis of codon optimized (for *E. coli* expression) DNA of the depicted organism. Level (B) consists of single expression modules into Dual expression vectors (Duet). Level (C) contains the dual gene expression modules vectors. Non-arrowed lines indicate the appropriately named restriction sites, while such lines with a red “X”, show respective restriction site deactivation. Letter symbols correspond to suitable Sequences: P_T7_, T7 Promoter; O, Lac Operator; T, Termination, E.c., *Escherichia coli*

All plasmid constructions used in this study are shown in Table 2 and Fig 2. The *TYR* gene was amplified from *Ralstonia solanacearum* genomic DNA with the primer pair RsTYRfw2/RsTYRrv2 (Table S1). The amplified DNA fragment was digested with the restriction enzymes *Nde*I and *Xho*I that flank the fragment and was inserted into the respective sites of an empty pRSF Duet vector (Novagen). The resulted vector was named pRSF-RsTYR (Fig 2). The *AADC* gene from *P. crispum (*parsley) was amplified from parsley root cDNA, where pathogen responsive genes [40] are usually active [41], with the primer pair PcTDC2bfw/PcTDC2bfw2 (Table S1). The amplified DNA fragment was digested with the restriction enzymes *Nco*I and *Bam*HI and inserted into the respective sites of the pRSF Duet vector generating the pRSF-PcTDC expression vector. To create a double gene expressing pRSF vector, the pRSF-RsTYR plasmid was digested with *Hin*dIII/*Xho*I enzymes to excise the RsTYR expression module and inserted into the similarly digested pRSF-PcTDC, thus creating the pRSF-PcTDC-RsTYR expression vector.

The translated sequence of RsTYR (497 aa) revealed a twin arginine translocase (TAT) signal peptide (1-34 aa) and a domain containing the two TYR characteristic copper-binding motifs (CuA, 93-100aa and CuB, 246-257 aa). The presence of TAT at the N-terminus suggests that the protein is exported to the periplasmic space of *R. solanacearum* as happens also with other tyrosinases (e.g. of the genus *Streptomyces*) that are also secreted via the TAT secretion pathway [42]. On the other hand, the translated sequence of PcTDC (515 aa) revealed a characteristic domain for AADC (data not shown).

Alternative forms of AADCs encompassing only a DOPA decarboxylase activity (DDC) were sought from *Aedes aegypti (*dihydroxyphenylacetaldehyde synthase, DHPAAS) or from *Sus scrofa (SsDDCmut*) and their sequences were inserted into pRSF and pCDF expression vectors (Table 1, Table S1). The DHPAAS coding sequence was codon optimized (named *AeDHPAASopt*) and synthetically constructed (GenScript, USA) for expression in *E. coli*. The appropriately designed 5’ and 3’ ends of the DHPAAS coding sequence (Genbank nucleotide sequence KT334547 (Table 2) was digested with the *Nco*I and *Hin*dIII restriction enzymes and inserted into the pRSF and pCDF vectors. On the other hand, a mutated version of the DDC gene (*SsDDCmut*) bearing an Y332F mutation was subcloned from the pKK223-3 (Bertoldi et al. 2002) to the pRSF and pCDF expression vectors to the *Nco*I/*Hin*dIII restriction sites. Utilizing a splice by overlap extension (SOE) PCR protocol [43] with the SsDDC1/SsDDCmut1 and SsDDCmut2/SsDDC2 primer pairs the inactivation of an internal *Nde*I restriction site was performed. The resulted vectors were named pRSF-AeDHPAASopt or pCDF-AeDHPAASopt and pRSF-SsDDCmut or pCDF-SsDDCmut respectively (Table 2).

#### Construction of hydroxytyrosol pathway expression vectors with modified genes

Two expression vectors with modified versions of TYR and AADC were constructed, based on the pRSF Duet vector. The modifications were designed to improve the co-localization of the two enzymes. The modified version of *P. crispum* TDC was created with splicing by overlap extension (SOE) PCR [43] in order to deliver a new species of AADC with a TAT signal peptide (TorA) attached to the N-terminal (Thomas et al. 2001) of the AADC for its translocation to the periplasm. The *TorA* and the *AADC* fragment without the translation initiation codon (ATG) were individually amplified (Table S1) and fused to create the coding sequence for the TorA:PcTDC enzyme (PcTDCmod, Genbank nucleotide sequence KT334545). The *PcTDCmod* was digested with *Nco*I/*Bam*HI in order to insert into the pRSF Duet at the respective sites generating the pRSF-PcTDCmod.

According to PRED-TAT algorithm [39] and the integrated web tools (http://www.compgen.org/tools/PRED-TAT/submit), the N-terminal of *R. solanacearum* TYR bears a peptide for translocation of the protein to the periplasm of *E. coli (*Fig S1) with reliability score of 0.981 (max value 1). Primers were designed (Table S1) to amplify the part of TYR CDS without the nucleotides responsible for the above-mentioned signal peptide carefully reconstituting the translation initiation codon (ATG). The amplification product that was 90 bp smaller than the wild type, was digested with *Nde*I/*Xho*I and inserted into the similarly digested pRSF Duet vector, thus creating the pRSF-RsTYRmod expression vector.

For the construction of the dual expression pRSF-PcTDCmod-RsTYR the pRSF-RsTYRmod was digested with *Hin*dIII/*Xho*I restriction enzymes and the RsTYRmod restriction fragment was inserted into the respective sites of pRSF-PcTDCmod. Similarly, the pRSF-PcTDC-RsTYRmod vector was constructed by the insertion of the *Hin*dIII/*Xho*I restriction fragment of pRSF-RsTYRmod into the respective sites of pRSF-PcTDC.

#### Construction of aldehyde reductase expression vectors

Two aldehyde reductase (ALR) genes from *E. coli* BL21(DE3) (*yqhD*, and *yahK*) were cloned into pRSF Duet and pCDF Duet vectors. The *yqhD* and *yahK* ALRs were amplified with the yqhD-fw1/yqhD-rv1 and *yahK*-fw1/*yahK*-rv1 primer pairs respectively (Table S1) and were cloned into pCDF Duet vectors after digestion of both PCR and vector with *Nco*I/*Pst*I. The resulted plasmids were named pCDF-yqhD or pCDF-yahK respectively Table S1.

The phenylacetaldehyde reductase gene (*PAR*) from *Rhodococcus erythropolis (*RePAR) was amplified with the primer pair RePARfw2/RePARrv2 (Table S1). The amplified DNA fragment was digested with the restriction enzymes *Nco*I and *Hin*dIII and was inserted into the respective sites of an empty pET Duet vector resulting the pET-RePAR vector (Table 2)

#### Construction of monoamine oxidase and phenylacetaldehyde reductase expression vectors

The monoamine oxidase gene from *Micrococcus luteus (MlMAO*) was amplified with the MlMAOfw7/MlMAOrv7 primer pair. The amplified DNA fragment was digested with the restriction enzymes *Nde*I and *Eco*RV and inserted into the respective sites of an empty pET Duet vector as well as in the pET-RePAR vector generating the pET-MlMAO and pET-RePAR-MlMAO expression vector respectively (Table 2).

#### *In-vivo* expression and growth optimization experiments

All genes referred in Table 1 were used primarily in *in-vivo* experiments, in order to check their optimal expression as well as their function. The latter was achieved by measuring their activity of their respective enzyme product activity through the quantity of their equivalent end-product biosynthesis. Two growth protocols were used in this work, targeting the maximal production of each intermediate step as well as the corresponding end-product biosynthesis; the LB-M9 and the M9 protocols. The LB-M9 protocol was utilized in feeding experiments while the M9 protocol was utilized for HT production directly from glucose.

#### Protocol LB-M9

A 5 ml starter culture was set in LB (1 % tryptone, 0.5 % yeast extract, 1 % NaCl) from freshly transformed *Escherichia coli* with appropriate plasmids and left to grow overnight at 37°C. The next day, 50 ml of LB were inoculated with the overnight culture to OD_600_=0.05-0.1 and were left to grow to an OD_600_=0.4-0.6. At that point Isopropyl β-D-1-thiogalactopyranoside (IPTG) was added at the selected concentration to induce the expression of the heterologous gene(s). After an induction time of 3 hours the cells were harvested by centrifugation (6000 g, 10 min), suspended in M9 medium (238.7 mM Na_2_HPO4·7H_2_O, 110.2 mM KH_2_PO_4_, 42.8 mM NaCl, 93.5 mM NH_4_Cl, 50 mM glucose, 1 mM MgSO_4_, 0.1 mM CaCl_2_, 10 nM thiamine) and appropriate substrate was added. Fermentations were performed in 250 mL Erlenmeyer flasks with orbital shaking at 200 rpm and 30°C. For the analysis of the substrate consumption and the product biosynthesis, 1 ml samplings were collected and kept at − 20°C until their electro-chromatographic analysis (see below).

#### Protocol M9

Similarly to LB-M9, a 5 ml starter culture was set in LB with appropriate plasmids and left to grow overnight at 37°C. The next day, 50 ml of LB were inoculated with the overnight culture to OD_600_=0.05-0.1 and were left to grow to an OD_600_=0.4-0.6. The cells were harvested by centrifugation (6000 g, 10 min), washed with M9 broth containing the basic salts (238.7 mM Na_2_HPO4·7H_2_O, 110.2 mM KH_2_PO_4_, 42.8 mM NaCl, 93.5 mM NH_4_Cl) and resuspended in M9 broth as in LB-M9 protocol. For the protein induction IPTG was added at the selected concentration and appropriate substrate was added. Fermentations were performed in 250 mL Erlenmeyer flasks with orbital shaking at 200 rpm and 30°C. For the analysis, 1 ml samplings were collected and kept at −20°C until their electro-chromatographic analysis.

### Tyrosinase and aromatic amino acid decarboxylase *in-vitro* enzymatic determination

To obtain bacterial cellular extracts enriched in the expressed proteins, freshly transformed BL21 clones with the appropriate expression vector were grown at 37°C overnight in 5 ml LB. The next day 50 ml of LB in a 250 ml Erlenmeyer flask were inoculated with 1 ml of the overnight culture and left to grow for 3 hours to reach and OD_600_ of about 0,5. At that point the inducer IPTG was added to a final concentration of 1 mM. Cells were left to grow for 20 hours and then the cells were harvested by centrifugation (3000 g, 10 min, 4°C) and washed with 50 ml of cold 20 mM Tris·HCl pH 7, harvested again and resuspended in 5 ml of the same buffer. The cells were disrupted by sonication with a Braun Labsonic U sonicator (5-min treatment at a relative output power of 0.5 with 0.5 duty period). The homogenate was centrifuged at 20000g for 30 min, and the supernatants were used for enzymatic activity determinations.

TYR activity was determined as previously described [44] for tyrosine hydroxylase. Briefly, TYR activity was carried out at room temperature in an assay (1 ml) containing 0.05% SDS, 20 mM phosphoric buffer pH 5, 1 mM tyrosine, 25 μM CuSO_4_. The reaction was initiated upon the addition of a crude extract containing 500 μgr of total proteins (described above) estimated with the Bradford assay [45]. Successive samplings of 100 μl at 0, 10, 20, 30, and 60 minutes after reaction start were analyzed in CE.

AADC activity was determined as previously described [27]. Briefly, the reaction mixture (1 mL) contained 1 mM aromatic amino (tyrosine or DOPA), 50 μM pyridoxal-5’-phosphate (PLP), 1 mM ascorbic acid in 100 mM Tris·HCl (pH 7.2). The reaction was initiated upon the addition of a crude extract containing 500 μgr of total proteins (described above) estimated with the Bradford assay [45]. The reaction was started by addition of the enzyme solutions and carried out at room temperature. Successive samplings of 100 μl at 0, 10, 20, 30, and 60 minutes after reaction start were analyzed.

### Electro-chromatographic analysis

Qualitative and quantitative analysis of samples were carried out in an Agilent Capillary Electrophoresis (CE) G1605 system coupled with a Diode Array Detector (DAD). A bare fused silica capillary column was used with effective length of 50 cm and inner diameter of 75 μm. The analysis was carried out in three steps: preconditioning, inlet and outlet buffer replenishment, buffer customization, injection and analysis. The preconditioning step involved an initial flush with 1 N sodium hydroxide for 2 min followed by a flush with analysis buffer (25 mM sodium tetraborate decahydrate, pH 9.2) for 4 min. Buffer customization includes the application of 20 kV voltage for 2 min before each analysis The injection step comprised a 50 mbar pressure for 2 sec and the analysis was performed at 29 kV for 10 min, time sufficient for maximum resolution of the compounds analyzed in this study. The temperature of the column cassette was kept constantly at 25°C. Replenishment and buffer customization steps were necessary to eliminate poor reproducibility of the peaks resulted from electrolytic phenomena in the running buffer and compound instability. Compounds under study were identified by matching the retention time, UV-absorbance spectrum, and co-chromatography with authentic chemicals. Calibration curves were obtained with authentic compound solutions of various concentrations. All samples were in triplicates unless stated otherwise.

### Mass spectrometry analysis of HT production

Further qualitative and quantitative analysis was performed to the most efficient metabolically engineered *E. coli* strains that produced the highest concentration of HT. Those *E. coli* strains (HMS174-HT^OP^-yahK, see Table 2) were grown in 5L flasks filled with 1L of growth medium. After 48h growth, samples of metabolically engineered *E. coli* strains showing highest HT production were further analyzed through mass spectrometry.

For the qualitative and quantitative monitoring of the *E. coli* strains an Accela HPLC system (Thermo), coupled to a hybrid LTQ Orbitrap Discovery XL mass spectrometer (LC–HRMS) and equipped with an electrospray ionization source (ESI) was used. The separation was conducted using a gradient elution system consisted of water with 0.1% (v/v) formic acid (A) and acetonitrile (B). The elution started with 5% B and reached 50% in 5 min. After 2 min the system returned to the initial conditions and stayed for 5 min for equilibrium of the column. The flow rate was set at 0.4 ml/min and the total run time was 15 min. For the chromatographic separation a Fortis C-18 (1.7 μm, 150 x 2.1 mm) column was used heated at 40°C. The injection volume was 10 μl and samples were maintained at 10°C during analysis. Ionization was achieved in negative ion mode (ESI-) at 350°C. The mass spectrometric parameters were: sheath gas and aux gas flow rate 40 and 10 units respectively; capillary voltage −40 V and tube lens −69 V. The mass range was adjusted from 113 to 1000 *m/z*.

In order to quantify HT, calibration curves were built using six different concentration levels of the analyte. The selected levels were 1 μg/ml, 2 μg/ml, 4 μg/ml, 6 μg/ml, 8 μg/ml and 10 μg/ml. 2.4-Dinitrophenol was used as internal standard (IS) in the concentration of 0.3 μg/ml. The construction of the calibration curve based on the ratio area of HT/IS versus the concentration of HT. Linearity was evaluated by coefficient of determination (R^2^) from the linear regression using the least squares equation. All concentration levels were measured in triplicates. Blank samples were injected every three injections in order to avoid carryover effect of the different concertation levels. Data were acquired, analyzed and processed with the Thermo Xcalibur 2.1 software.

For the bacterial strain extraction, 225 mL of cell culture were centrifuged at 3500 rpm for 20 min. The supernatant was extracted with the adsorption resin Amberlite XAD7 after overnight treatment, in order to obtain an extract being enriched in phenolics. Methanol was used as extraction solvent. The obtained methanol extract was dried under vacuum until dry and reconstituted to 100 μg/ml for the HPLC-ESI-HRMS analysis.

## Results

### Tyrosine overproducing strain

Two plasmids/modules harboring 11 genes in total to produce tyrosine from phosphoenolpyruvic acid (PEP) and erythrose-4-phosphate (E4P) were created [29] by the utilization of the BglBrick system [46] and the relative transformants were behaving as tyrosine overproducers (OP strains, Table 2). Six of these genes were necessary for the biosynthesis of shikimic acid and five of them required for the conversion of shikimic acid into tyrosine. The shikimic acid biosynthesis module (pS4; Fig S2) bears the following genes under the lacUV5 or tetO promoters: phosphoenolpyruvate synthase (*PpsA*), transketolase A (*TktA*), 3-deoxy-D-arabino-heptulosonate synthase (*AroG*), Dehydroquinate synthase (*AroB*), Dehydroquinate dehydratase (*AroD*), shikimate dehydrogenase (*AroE*). The tyrosine module (pY3; Fig S2) contains genes under the lacUV5 or the trc promoters: chorismate mutase/prephenate dehydrogenase (*TyrA*), tyrosine aminotransferase (*TyrB*), chorismate synthase (*AroC*), 5-enolpyruvoylshikimate 3-phosphate synthase (*AroA*), shikimate kinase II (*AroL*). The 2 plasmids (pS4 and pY3) bearing all the necessary genes were inserted into BL21 and HMS174 *E. coli* strains (Table by electroporation. The resulted tyrosine-overproducing strains were evaluated for their ability to produce tyrosine under various conditions [29].

### Expression of genes involved in the biosynthesis of Hydroxytyrosol

#### *Ralstonia solanacearum* tyrosinase

The hydroxylation of the phenolic ring of tyrosine or tyrosol to DOPA or HT may be catalyzed by the action of the *R. solanacearum* TYR (Table 1). BL21 cells bearing the RsTYR (BL21-pRSF-RsTYR, Table 2), in induced conditions were checked initially for their ability to induce the expression of the TYR protein (Fig S3a). As negative controls BL21 cells were used transformed either with empty pRSF vector in non-induced conditions or with BL21-pRSF-RsTYR in non-induced conditions. It was obvious that a band with much higher intensity (see arrow in Fig S3a) than the non-induced or the empty vector treatments, was present above 43 KDa, corresponding to the TYR protein size of about 50 kDa [36]. Crude extracts from these cultures (non-induced BL21-pRSF, non-induced BL21-pRSF-RsTYR, and induced BL21-pRSF-TYR) were tested for TYR activity in a colorimetric assay since the dual activity of the enzyme oxidizes DOPA to the orange dopachrome and eventually to blackish melanins (Fig S3a). Conclusively, both the extracts from the induced and non-induced conditions were able to transform tyrosine to DOPA (and eventually to melanins) indicating from one hand the activity of the enzyme and on the other hand the leaky expression of TYR in non-induced conditions.

Subsequently, TYR activity was verified by *in-vitro (*Fig 3a) as well as colorimetric assays (Fig 3b). A time course experiment following tyrosine and DOPA certified that tyrosine was promptly converted to DOPA (Fig 3a). In all cases the color of the reaction was getting black indicating the melanization of DOPA through the diphenolase activity of TYR.

**Fig 3.**
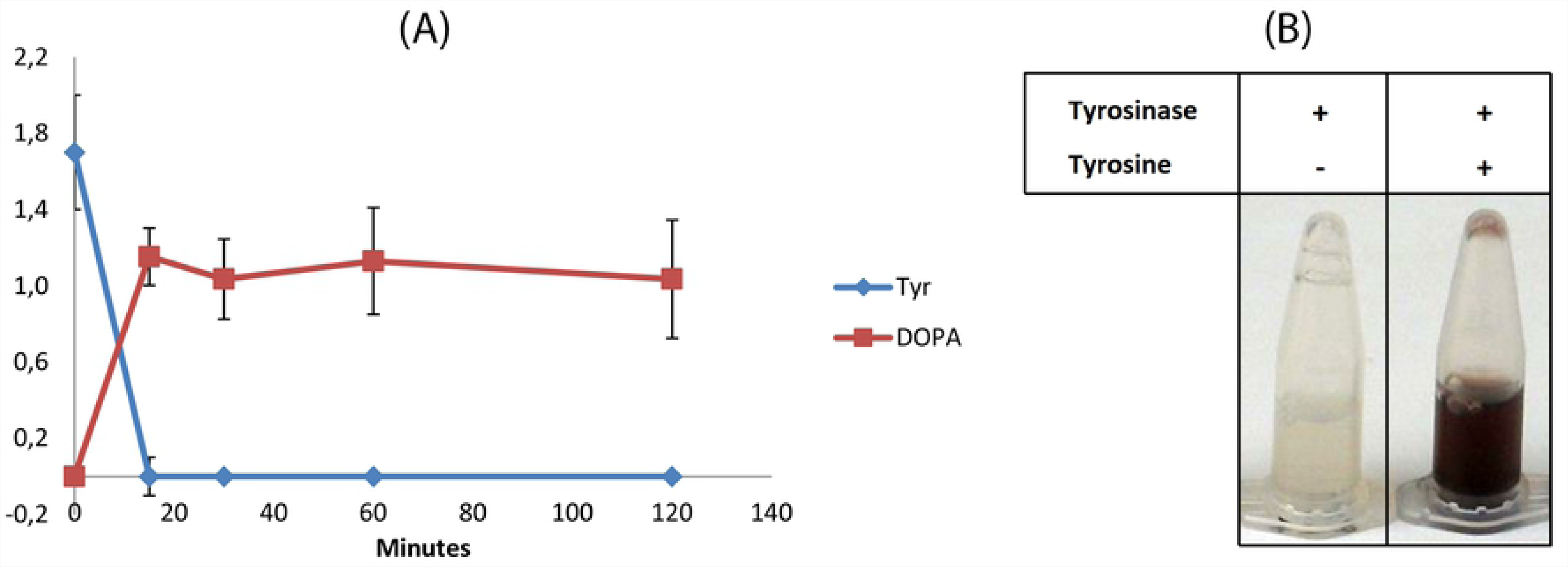
*In vitro* assays for the evaluation of tyrosinase activity. (A), The reaction was supplemented with 2 mM of tyrosine and DOPA production was followed. (B), Evaluation of initial tyrosine concentration on the activity of tyrosinase. (C), Blackening of reaction due to the melanization of the produced DOPA (o-diphenolase activity of RsTYR).

Furthermore, *in-vivo* assay for the conversion of 2 mM tyrosine or tyrosol to DOPA or HT respectively was set up (Fig 4). In induced conditions, the substrate (tyrosine or tyrosol) was converted to DOPA or HT respectively (Fig 4 D) 2h after the addition of the substrate. The decrease in DOPA concentration over time (Fig 4 B) is attributed to the sensitivity of the DOPA in the specific conditions of the culture. When the cultures were kept under non-induced conditions no consumption of the substrate was recorded or the consumption was minimal. A slight increase of the products was also recorded (Fig 4 A and C).

**Fig 4.**
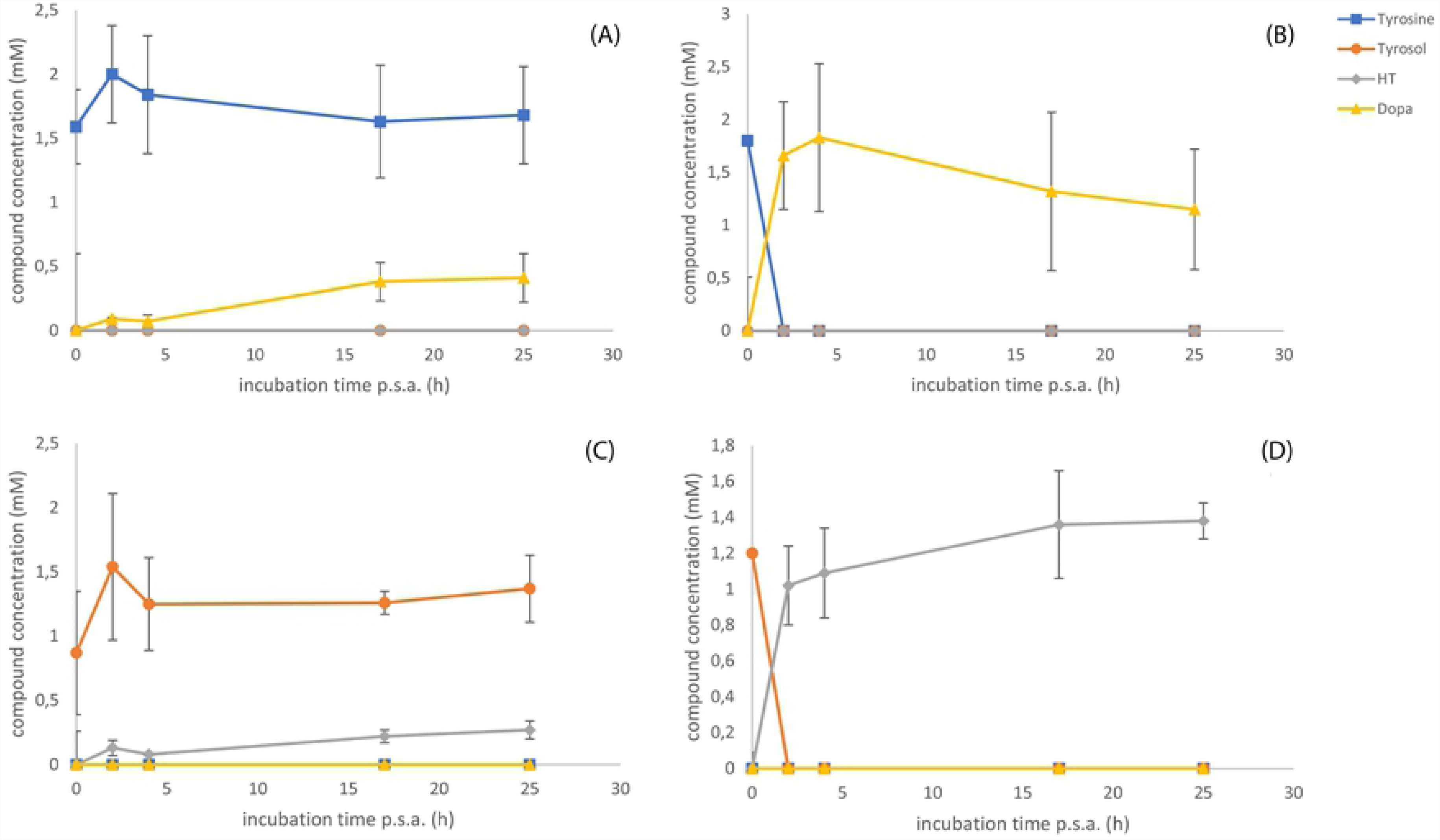
A. Protein expression in *Escherichia coli* to certify the tyrosinase protein expression. In the first lane total proteins from *E. coli* BL21(DE3)-pRSF in non-induced conditions were loaded. In the second, total proteins from BL21(DE3)-pRSF-RsTYR in non-induced conditions were loaded while in the third lane total proteins from BL21(DE3)-pRSF-RsTYR in induced conditions were loaded. The arrow in the protein marker helps to estimate the size of the protein band. B. Colorimetric assay with the protein crude extracts to assess the activity of tyrosinase as described in Material and Methods.

#### *Petroselinum crispum* aromatic amino acid decarboxylase

Beyond the hydroxylation of the phenolic ring, another essential reaction is the decarboxylation-amination of the aromatic amino acid by the respective AADC. Newly transformed BL21 cells bearing the PcTDC (BL21-pRSF-PcTDC, Table 2) in induced conditions were checked for their ability to induce the expression of the AADC (Fig S3B). As negative controls BL21 cells were used transformed either with empty pRSF vector in non-induced conditions or with BL21-pRSF-PcTDC in non-induced conditions. As with the case of TYR producing BL21 cells, a band with much higher intensity than the non-induced or the empty vector treatments, was present at about 50 KDa [40], corresponding to the AADC protein (Fig S3B).

*In-vivo* assays for the evaluation of AADC were set up. Induced cultures were supplemented either with tyrosine or DOPA and time course samplings were analyzed in CE (Fig 5). Either substrate tyrosine or DOPA was converted to tyrosol or HT respectively (see Fig 1 for biosynthetic scheme), with the latter substrate (DOPA) to have been consumed at about 5 hours after substrate addition. Similarly to TYR *in-vivo* assays, the cultures under non-induced conditions presented minimal consumption of the substrate (Fig 5 A and C) while the induced conditions exerted the maximal conversion efficiency (Fig 5 B and D).

**Fig 5.**
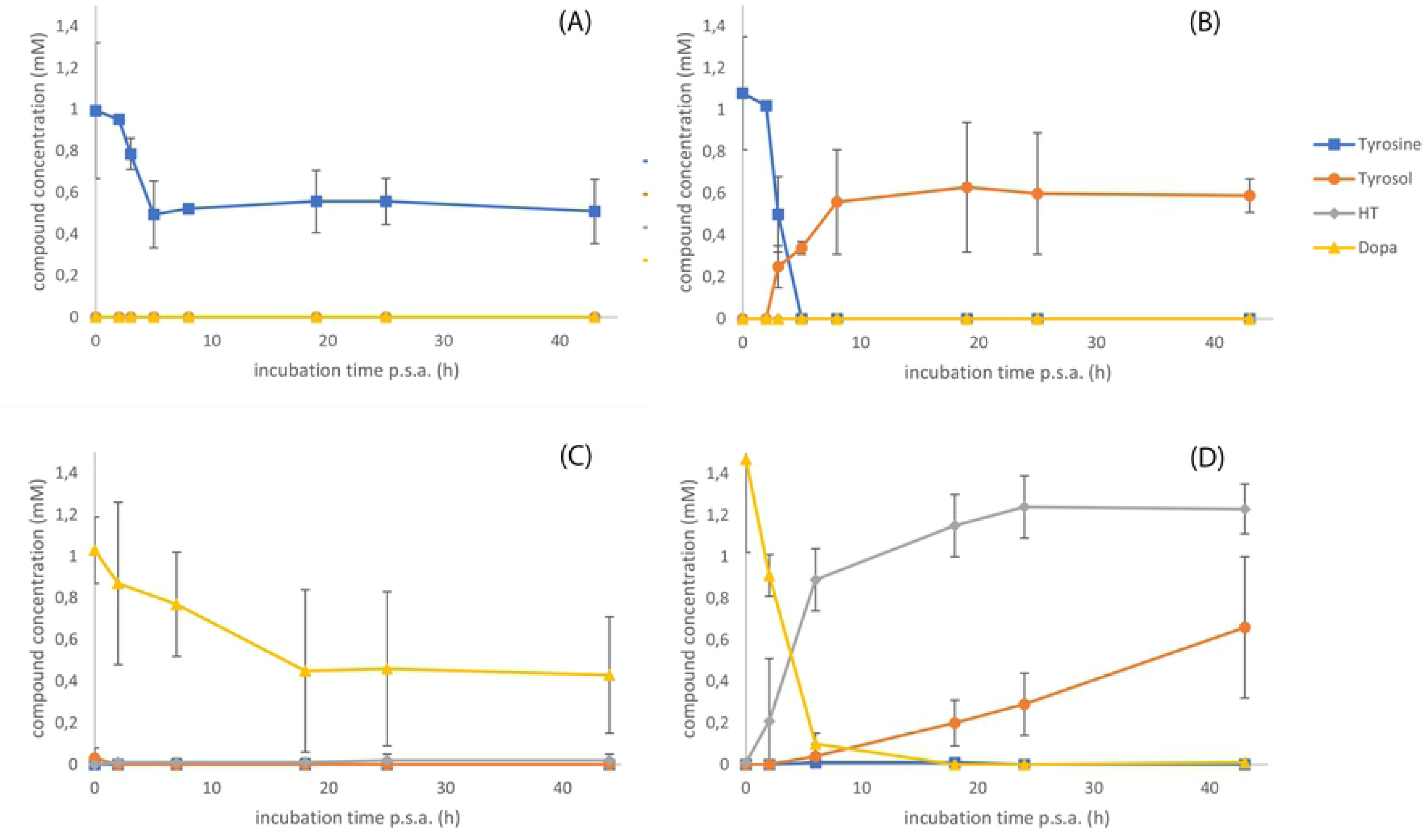
*In-vivo Petroselinum crispum* aromatic amino acid decarboxylase assay acting on supplemented tyrosine or DOPA. Panes A and B refer to the *in-vivo* experiment supplied with tyrosine while panes C and D is referred to the experiment supplied with DOPA. The panes at the left of the figure (panes A and C) depict non-induced conditions while the panes on the right part depict (panes B and D) the experiment from induced conditions.

### The expression of MlMAO, RePAR, AeDHPAASopt, SsDDCmut, PcTDCmod or RsTYRmod did not benefit HT production strategy

Until we come to the basic transcription units for the production of HT (pDUET vector bearing RsTYR and PcTDC genes, Fig 2), several different constructs were evaluated utilizing diverse combinations of transcription units in order to find the optimal approach that would lead to the maximal production of HT. Although, all the coding sequences of *MlMAO*, *RePAR*, *AeDHPAASopt*, *SsDDCmut*, *PcTDCmod* or *RsTYRmod (*Table 1, Table 2, Table S1) under the IPTG-inducible T7 promoter were evaluated on different media, different induction temperatures, different induction protocols, different vectors with different origin of replication, no specific advantage was gained over the basic PcTDC-RsTYR pair (detailed data are not shown). In all these cases, *in-vivo* expression experiments were set-up, heterologous proteins were induced, and samples were analyzed to estimate their activity.

### HT production from engineered *E. coli* strains

The proof that *RsTYR* and *PcTDC* genes were active, was followed by the combined *in-vivo* expression. Cultures of BL21 cells transformed with pRSF-PcTDC-RsTYR (Table 2) were set up to evaluate their ability to convert supplemented tyrosine into HT in both induced and non-induced conditions (Fig 6A). As a rule, 1 mM of tyrosine was supplemented because no worthwhile differences on the production of HT were observed when higher concentrations were evaluated (data not shown). When no tyrosine was supplemented, 0.21 mM of HT (Fig 6C) was produced directly from glucose (at 48h of cultivation).

**Fig 6.**
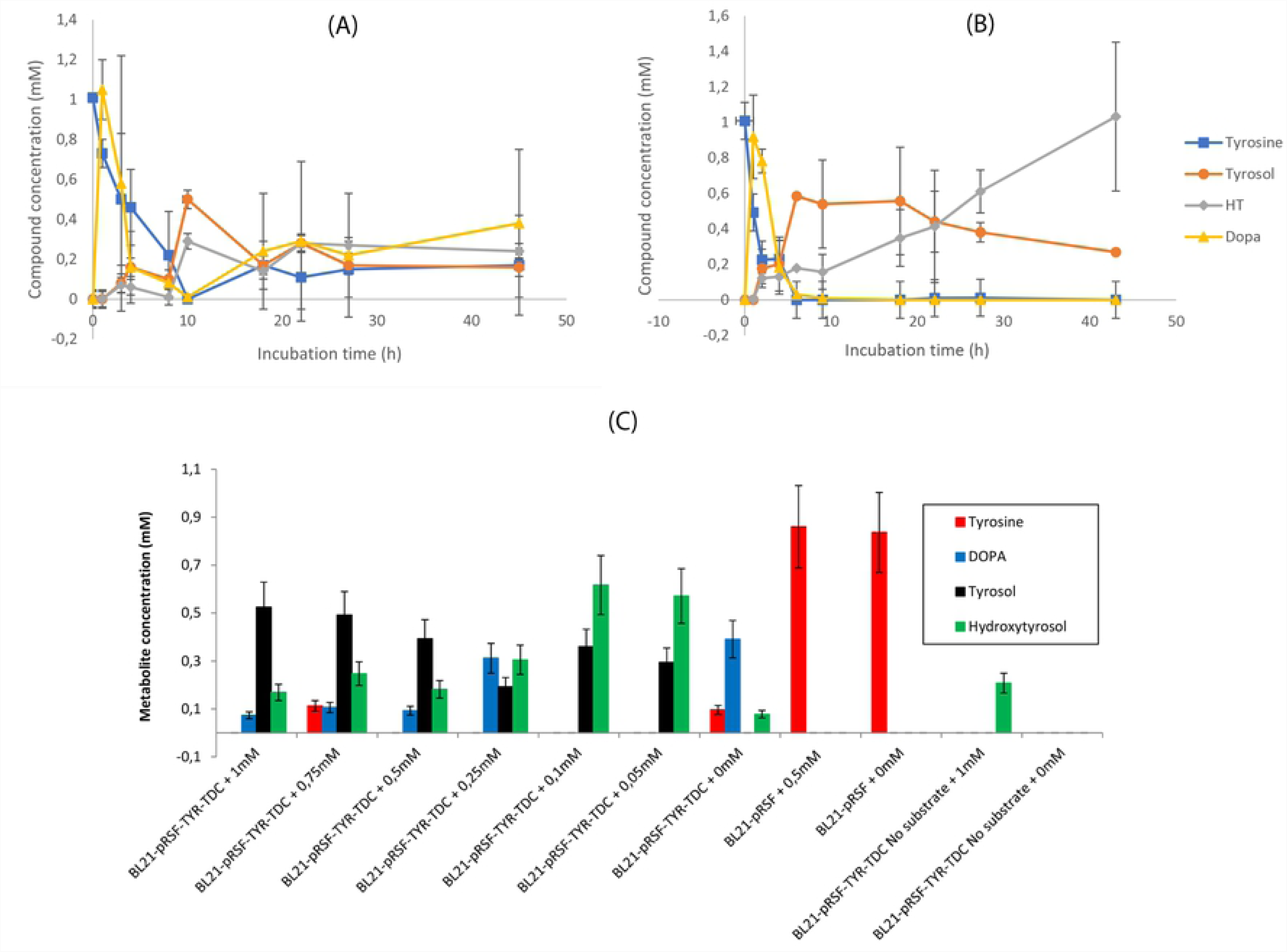
*In-vivo* evaluation of the BL21-TYR-TDC for its ability to produce hydroxytyrosol (HT). (A) non-induced conditions. (B), induced conditions. Tyrosine was initially supplemented at 1mM concentration and HT production as well as the intermediate compounds, DOPA and Tyrosol, were estimated. (C) Evaluation of IPTG concentration on hydroxytyrosol production from the BL21-pRSF-TYR-TDC strain (Table 2). Values are from 48 h post substrate addition.

From the observation of non-induced conditions (Fig 6A) it is shown that there is activity of both the TYR and AADC advocated by the presence of DOPA or tyrosol although in minimum levels. On the other hand, the chart presenting the induced conditions (Fig 6B) shows that most of the supplemented tyrosine (about 80%) was rapidly converted into DOPA that rapidly disappeared within the next 6 h. Tyrosol appears too, as early as DOPA, although at low concentrations and then is stably increased to a maximum of about 0.6 mM as soon as 7-8 h of incubation. HT appears at the same time with tyrosol, but up to 10h of incubation remains almost at the same levels. When tyrosol starts decreasing after about 18h of culture incubation, HT shows a second phase increment reaching a maximum of about 1 mM.

### Effect of IPTG concentration on hydroxytyrosol production

Experiment with protein induction occurring in increasing concentrations of IPTG was evaluated in order to find the ideal combination of protein expression-enzyme activity. The evaluated strain was the BL21-pRSF-TYR-TDC (Table 2) supplemented with 1mM of tyrosine and the evaluated IPTG concentrations were 0, 50, 100, 250, 500, 750, and 1000 μM. In all cases the system activity was observed through the analysis of the final as well as the intermediate compounds tyrosine, DOPA, tyrosol, and HT (Fig 6C). Both TYR and AADC were active suggested by the presence of DOPA, tyrosol or HT in the medium. A first note is that there is substrate loss i.e. to melanin accumulation through the oxidation of the tyrosine-derived DOPA through dopachrome. The rest of tyrosine was intracellularly converted into tyrosol or HT by the action of the AADC or the AADC and TYR respectively. As IPTG concentration was decreased from 1 mM to 0 mM, differences were observed in the produced analytes; the intermediate tyrosol is lowering while the concentration of the final product HT is increased. Furthermore, in the lower IPTG concentrations (50 and 100 μM) DOPA is not traced. These advocate to a more balanced system for the production of HT. When no tyrosine was added to the system a minimal amount of HT was produced in induced conditions from glucose or no HT was produced in non-induced conditions. The negative control BL21 cells that did not carry the HT pathway genes and did not consume the exogenous supplied tyrosine. Consequently, 50 μM of IPTG was regarded as the optimal concentration of the inducer and was used throughout our experimental conditions unless stated otherwise.

### Hydroxytyrosol production from engineered tyrosine-overproducer *E. coli* strains

Instead of using an *E. coli* system that is necessary to be supplemented with the precursor amino acids tyrosine or DOPA to produce HT, a biological system designed to overproduce tyrosine was further engineered by introducing the pS4 and pY3 plasmids to BL21 and HMS174 hosts, for the production of HT directly from glucose (thus named BL21-HT^OP^ or HMS174-HT^OP^, Table 2). However, preliminary experiments with BL21-HT^OP^ and HMS174-HT^OP^ showed non-consistent tyrosine production, while HT levels remained similar reaching the level of 0.21mM, when both strains evaluated on M9 medium (Fig S5). This was much an unexpected result, since there was a noticeable difference on tyrosine pool, between the two strains. In HMS174-HT^OP^ tyrosine levels reached 0.63mM, while in BL21-HT^OP^ appeared that all internally overproduced tyrosine was at nearly non-existing levels indicating that it was all consumed in primary metabolism. Similar biosynthetic equimolar levels were obtained for the intermediate metabolites DOPA and tyrosol in HMS174-HT^OP^, however, leading to still low HT level. This metabolic event indicated that a probable biosynthetic bottleneck existed in the direct sequential conversion of the internally produced tyrosine to hydroxytyrosol.

#### Auxiliary aldehyde reductase genes

The biosynthetic pathway of this work started by utilizing the dual specificity of AADC from *P. crispum (*Table 1) with respect to the amino acid that accepts as substrate. It is able to convert tyrosine or DOPA to 4-hydroxyphenylacetaldehyde (HPAA, monophenol route) or 3,4-dihydroxyphenylacetaldehyde (DHPAA, diphenol route) respectively (Fig 1). The next step should be the reduction of the phenylacetaldehyde to the corresponding alcohol, tyrosol or HT (Table 1, Fig 1). *E. coli* is an organism that possesses several ALRs that drive the previously mentioned conversion [27]. We therefore checked whether the presence of extra overexpressed ALRs could confer any advantage to the engineered system. The ALRs evaluated were the YahK (ALR-K), and the YqhD (ALR-D) [30] both of which are present in the genomes of both BL21 and HMS174. However, while the over-expression of *yqhD* along with the HT biosynthesis genes did not give any advantage to the production system (data not shown), this was not the case for the over-expression of *yahK*. While the over-expression of the *yahK* gene resulting in BL21-HT^OP^-yahK strain (Table 1), caused a 62% decrease in the production of HT, it turned to 386% increase when the host was the HMS174 (HMS174-HT^OP^-yahK, Table 1), as shown in Fig 7a reaching 1.02 mM (157.2 mg/L), when analyzed in CE-DAD system. All the follow up experiments continued with optimization of only the highest HT titer producing strain, HMS174-HT^OP^-yahK.

**Fig 7.**
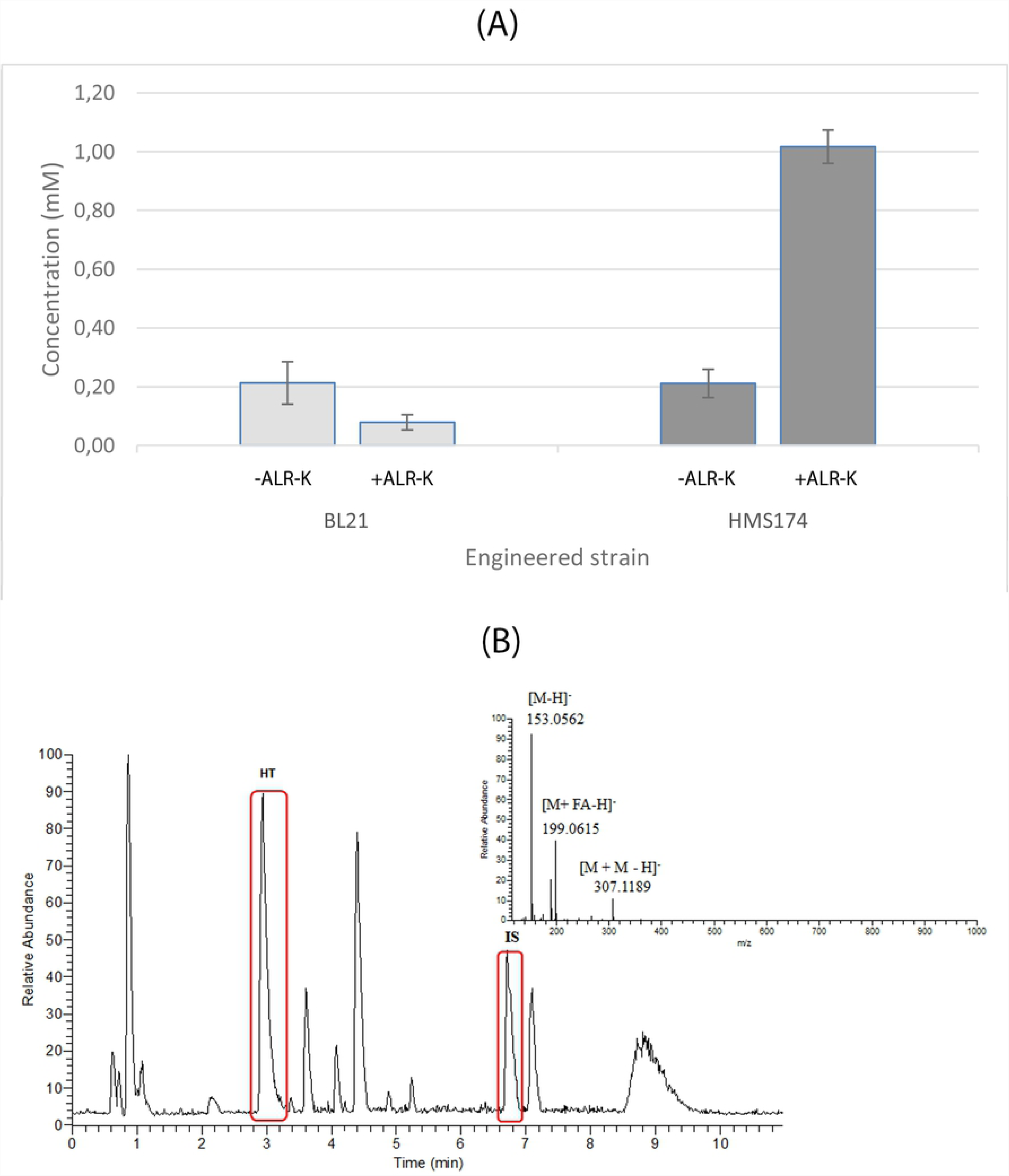
Evaluation of hydroxytyrosol production from BL21-HT^OP^ or HMS174-HT^OP^. (A) Evaluation of the effect of *yahK* aldehyde reductase gene (*ALR-K*) on the production of hydroxytyrosol when BL21-HT^OP^ or HMS174-HT^OP^ strains were used as hosts. The values were extracted at 48 hours of cultivation. The effect of the *yahK* in the HT production is also presented as a percentage value (negative or positive) above the bar representing the strain engineered to express the *yahK* gene. (B) Base-peak HPLC-ESI-MS chromatogram of the methanol extract of the strain HMS174-HT^OP^- yahK. The two peaks circled in red are Hydroxytyrosol (HT; left) and Internal Standard (IS; right). At the mass spectrum the characteristic ions of HT are denoted (pseudomolecular ion [M-H]^-^ of HT at *m/z* 153.0562, the [M+FA-H]^-^ adduct ion of HT with formic acid and the pseudodimer ion [M+M-H]^-^ of HT), produced during the ionization process.

#### Optimal analysis of the highest HT producer strain through Mass Spec analysis

Furthermore, the strain HMS174-HT^OP^-yahK (Table 2), carrying the full HT overproducing pathway (involving overexpressed genes, *RsTYR*, *PcTDC* and *ALR-K / yahK*, Table 1), as well as all the genetic accessory to overproduce tyrosine ([29], Table 1), was further analyzed by a LC-MS-MS mass spectrometry. In this highest titer strain the precursor of HT pathway, tyrosine (Fig 1) was internally produced by feeding the strain with glucose supplemented to the growth medium. This strain grown under the protocol M9, showed the highest HT production when samples were analyzed from 50ml cultures (Fig 7A) with a CE-DAD system. However, it was important to show whether there was any effect at HT production on large growth medium volume and analyzed by mass spectroscopy. Indeed, the MS analysis showed HT production at much higher levels.

After the cleaning and enrichment process of the strain HMS174-HT^OP^-ALR-K using adsorption resins, the dried weigh of the methanol extract was estimated at 4.2 mg/ml cell culture. In order to determine the HT levels HPLC-ESI(-)-HRMS was employed. Fig 7b illustrates the obtained base-peak chromatogram of the extract after resin treatment together with the corresponding mass spectrum of HT. As it is highlighted in red, HT is eluted at 2.94 min and IS at 6.71 min. Other peaks are also detected mainly representing intermediate compounds of the metabolic pathway (*data not shown*). At the mass spectrum (Fig 7B) the ions corresponding to HT are presented. The high accuracy and resolving power of the Orbitrap analyzer as well as the use of reference standard enables the unambiguous identification of HT in the complex mixture [47]. Together with the pseudomolecular ion of HT ([M-H]^-^), other ions corresponding to its adducts with formic acid ([M+FA-H]^-^) and its dimer ([2M-H]^-^) were also detected. Using Thermo Xcalibur 2.1 software and extraction ion method (XIC), the quantification of HT took place using linear regression method. A standard calibration curve was built described by the equation y=0.1674x + 0.1929 while coefficient of determination was calculated R^2^=0.9932, indicating the good linearity achieved. Based on the equation, the levels of HT were calculated at 270.8 mg/L of cell culture.

## Discussion

In this project we succeeded to modularly engineer both the primary and secondary metabolism of *E. coli* to produce HT. We chose a rational approach for its production [16] where the primary metabolism was engineered in order to boost tyrosine production that is subsequently utilized for the biosynthesis of HT through the engineered secondary metabolic machinery.

The HT molecule bears a benzene (phenolic) ring linked with an alcoholic chain with two carbon atoms (Fig S4). Since, all phenolic compounds encountered in organisms emanate from the aromatic amino acids via the shikimic acid pathway [48, 49], we assumed that this is also the case for HT biosynthesis. From the three aromatic amino acids of the primary *E. coli* metabolism (phenylalanine, tyrosine, tryptophan), HT resembles most with tyrosine (Fig S4) and could be biosynthesized by a hydroxylation, a decarboxylation, a deamination, and finally a reduction (Fig 1B).

The first major biochemical conversion that had to be achieved was the hydroxylation of tyrosine leading to the formation of DOPA. Hydroxylation of tyrosine could also be performed by a pterin-dependent monooxygenase (tyrosine hydroxylase, E.C. 1.14.16.2) that specifically hydroxylates the phenolic ring [50]. Alternatively, hydroxylation could occur by a type III copper TYR (E.C. 1.14.18.1) that hydroxylates the phenolic ring of tyrosine, but as a feedback reaction it could also oxidize the hydroxyl groups to quinones in subsequent steps [24, 36]. Both metabolic engineering solutions retain pros and cons; the former could be considered as a probable choice for hydroxylating tyrosine to HT but needs the co-factor tetra-hydro-biopterin that is absent from bacteria [27]. Alternatively, the latter choice can “over-oxidize” the produced DOPA to the problematic ortho-quinone metabolites [51] resulting in an undesirable product for our target.

Thus, we chose to utilize the TYR from *R. solanacearum (*Table 1) for the following reasons. Although, tyrosinases show a much higher specific activity for the oxidation of o-diphenols (e.g. formation of DOPA quinone; diphenolase or o-diphenol oxidase activity) than for the hydroxylation of monophenols (e.g. DOPA formation; monophenolase or monophenol hydroxylase activity), the TYR from *R. solanacearum* possesses a higher monophenolase than o-diphenolase activity [36]. This difference can result in the redirection of the metabolic flow from tyrosine to DOPA (Fig 1B) with less toxic byproducts. The monophenolase activity of *R. solanacearum* TYR was confirmed in *in-vitro (*Fig as well as in *in-vivo (*Fig 4A and 4B) experiments on tyrosine as well as on tyrosol (Fig 4C and 4D).

The next biochemical conversion should be the decarboxylation of tyrosine or DOPA (produced by the action of TYR on tyrosine) resulting to the corresponding phenylacetaldehyde by the sequential action of an Aminoacid Decarboxylase (AADC) and a Monoamine Oxidase (Fig 1B). We choose to utilize an *AADC* gene from parsley [40] which translate to an enzyme that can accept either tyrosine or DOPA as substrates. The specified enzyme has also the advantage that beyond the basal decarboxylation reaction performs an extra sequential deamination step that leads to the production of the corresponding phenylacetaldehyde [33], thus it also possesses a phenylacetaldehyde synthase activity (PAAS). The activity of the parsley AADC (PcTDC) was confirmed in *in-vitro* and *in-vivo* experiments where reactions were supplemented with either tyrosine or DOPA (Fig 5).

The dual specificity of the utilized TYR (over tyrosine or tyrosol) and AADC (over tyrosine or DOPA) enzymes unwraps two possible metabolic routes for the production of HT; one through the decarboxylation-deamination of DOPA (diphenol route, Fig 1) for the production of DHPAA and one through the decarboxylation-deamination of tyrosine for the production of 4-hydroxyphenylacetaldehyde (monophenol route, Fig 1). Both parallel occurring metabolic routes, result in the same metabolic flow from tyrosine to the production of HT.

According to the proposed biosynthetic scheme, the last biosynthetic step for the production of HT should be the reduction of the produced aromatic phenylacetaldehyde (4-hydroxyphenylacetaldehyde produced from the monophenol route or DHPAA produced from the diphenol route) by an *ADH/ALR* gene. The main two strains used to host the heterologous dual pathway (BL21 and HMS174) are known to possess several ALR enzymes, that may support the reduction of the produced phenylacetaldehydes to the corresponding alcohols, tyrosol or HT respectively [27]. The simultaneous expression of *RsTYR* and *PcTDC* was evaluated by the ability of the system to transform the supplemented tyrosine or DOPA to the corresponding phenylacetaldehydes and then to the respective tyrosol or HT (Fig 6A and 6B). Conclusively, this TYR-AADC dual activity approach proved to be a beneficial strategy for the robust manufacturing of the high-valuable compound HT. Our ultimate target in this work was to achieve equimolar conversion of tyrosine, as a primary precursor, to HT as an end-product of the above-mentioned routes.

Analyzing the RsTYR *in-vitro* and *in-vivo* activity results (Fig 3 and Fig 4) it was found that the enzyme was active in both induced and non-induced conditions but more interestingly it performed a quick reaction, within the first 20 min with the *in-vitro* crude *E. coli* extract. This was advocated by the identification of DOPA in early time points upon the supplementation of tyrosine. The *in-vivo* rapid conversion could be explained by the assumption that RsTYR was transferred to the medium through the periplasmic space of the host; it has been shown that RsTYR possess a signal peptide for its transfer to the periplasmic space (Fig S1) thus explaining the appropriate activity to produce DOPA from tyrosine. The extracellularly-produced DOPA was not directly available to AADC, that was intracellularly expressed. This differential compartmentalization may explain the reason why extracellular DOPA could not be easily converted to HT and was wasted as melanized byproducts due to exposure of supplemented tyrosine to RsTYR, leading to the oxidation of DOPA accompanied by the blackening of the medium (Fig 3C). Consequently, the late increase in HT is assumed to be produced through the newly synthesized endogenous tyrosine and the monophenol route (Fig 1). In that case, the intracellularly AADC-synthesized tyrosol (accompanied with the action of an ALR) was transferred outside of the cell before its transformation to HT by TYR. The fact that in BL21-HT^OP^ cultures no DOPA was traced and no blackening of the medium occurred, was a proof of the above assumption (HT production not through diphenol route); the endogenously produced tyrosine was subjected to the activity of the AADC (and the activity of the ALR) for the production of tyrosol and eventually to its transformation into HT through the hydroxylation of tyrosol by the TYR (monophenol route, Fig 1).

If the abovementioned assumption was true, any attempt to co-localize the two enzymes should make a difference in the efficiency of the system. Thus, we tested two different constructs; the first contained the transcriptional unit of the TYR along with a modified TDC fused with a translocation to periplasm TAT signal peptide at its NH_2_-end [52]. The second construction contained the wild type parsley AADC with a modified TYR truncated at its NH_2_-end (TAT signal peptide removed) to prevent the translocation to the periplasm/medium. In both cases we wanted to demonstrate the effect of co-localization of the enzymes either in the periplasm/medium or in the cytoplasm respectively. However, both approaches were unsuccessful to improve the efficiency of HT production, by supplementing tyrosine (data not shown).

While the production of HT through the proposed system was successful, the presumed spatial separation of RsTYR and PcTDC was a drawback for the system; ideally the product of TYR should serve immediately as the substrate for the AADC. To overcome this bottleneck, we tried to include in our strategy the activity of two new genes; a codon optimized DDC for *E. coli* expression from the mosquito *Aedes aegypti (*AeDHPAASopt) or the DDC from *Sus scrofa (SsDDCmut*). Both these decarboxylases use the amino acid DOPA as the exclusive substrate and put through both the decarboxylation and deamination reactions (Fig 1) thus forcing the metabolism of tyrosine to follow only the diphenol route. Nevertheless, we could not get any activity from these genes in our system although many different expression conditions were tested including temperature, IPTG concentration, vector copy number, and host strain. Similar problem was highlighted by Achmon et al. [53] when they expressed in *E. coli* a *PAAS* from *Rosa hybrid*, a gene that is orthologous to *AeDHPAASopt* and *SsDDCmut*, indicating a general expression problem for these enzymes in *E. coli*. The solution they proposed was the expression of the *PAAS* gene in the pTYB21 vector (New England Biolabs), where an N-terminal fusion protein with the intein VMA1 from the *Saccharomyces cerevisiae* was created. We made similar fusions of *AeDHPAASopt* and *SsDDCmut* with VMA1 but did not get any activity (data not shown).

The ability of the engineered *E. coli* to bio-transform the supplemented precursor tyrosine into the valuable HT as well as the fact that tyrosine was the limiting factor for producing HT (since its production rate was rapidly reduced to zero after tyrosine consumption; Fig 5B), urged us to evaluate the case of producing HT from a tyrosine over-producing strain. The efficient biosynthesis of tyrosine was necessary to avoid the exogenous supplementation of tyrosine and thus making the biological production of HT more economically attractive. To this extend, *E. coli* strains containing the modules for a) the biosynthesis of shikimic acid (pS3), b) the production of tyrosine from shikimic acid (pY3), and c) the production of HT from tyrosine were utilized (BL21-HT^OP^ and HMS174-HT^OP^) for the production of HT. As was earlier mentioned, the supplementation of tyrosine into the medium of BL21-*RsTYR-PcTDC* or HMS174-*RsTYR-PcTDC* strains led to intermediate product formation due to spatial separation of the RsTYR and PcTDC activities. The channeling of intracellularly produced tyrosine to the monophenol route of HT pathway (Fig 1) resulted in the formation of tyrosol that was eventually excreted into the medium where it was converted to HT. However, both engineered strains resulted at this stage to non-equimolar levels of HT.

Further investigation revealed that HMS174-HT^OP^ was a much higher tyrosine overproducer than BL21-HT^OP^ (Fig S5). The fact that in the BL21-HT^OP^ no intermediate products were apparent opposite to HMS174-HT^OP^, where all previous to HT metabolites, tyrosine, DOPA and tyrosol, were not only in excess, but to non-equimolar levels, point to the assumption that a probable biosynthetic bottleneck exists that either consumes/converts an intermediate metabolite and/or possibly a new product affects a downstream biosynthetic step.

HT levels were still at levels below 1mM, indicating still non-equimolar favorable conditions. Thus, our work was further focused to the HMS174 strain that was the highest tyrosine overproducing strain. Further investigation revealed that involvement of an extra overexpressed gene from the *ADH*/*ALR* family of *E. coli* was able to lead to successful equimolar levels of HT produced from the HMS174-HT^OP^. Indeed, the question as to whether the activity of the native *ALR* genes could be the limiting factor in the HT production, was answered by over-expressing two *E. coli ALR* genes, the *yqhD (ALR-D*) and the *yahK (ALR-K*) along with the HT biosynthetic genes (Fig 1B) and the two plasmid gene cassettes (Fig 1A, [29]) responsible for the tyrosine overproduction. These two ALR genes were selected based on earlier evaluation of their ability to convert similar aldehydes into alcohols [30, 54]. Interestingly, while the over-expression of *ALR-D* did not cause any further improvement of HT production, the over-expression of *ALR-K* in HMS174-HT^OP^ strain increased the production of HT by 386% reaching a concentration of 1.02 mM (157.2 mg/L, Fig 7a) when analyzed by a CE-DAD, indicating a biosynthetic bottleneck was able to overcome by *ALR-K* over-expression.

Such a high increase in the HT production in the HMS174-HT^OP^ strain was, most likely, the result of the respective increase of the endogenous tyrosine pool, delivered by the overexpression of the total tyrosine biosynthesis pathway (11 overexpressed genes, Fig 1). Compared to the HMS174, the tyrosine pool in the BL21 strain was substantially lower (Fig S5) fact that explains the reduced HT titer in the latter strain (Fig 7A). However, further LC-HRMS analysis has shown an even higher HT production reaching 1.76 mM (270.8 mg/L, Fig 7B). This increase in HT production under these conditions was probably due to the stationary phase protein overproduction that is a fundamental capability of *E. coli* [55], which has been shown to lead to protein expression with higher protein solubility and thus higher enzyme activities [56]. Moreover, this biosynthetic incident reveals that ALR-K is relieving the pathway from the effect of a possibly deleterious accumulating product, over the previously tested biosynthetic scheme (with ALR-K absent or poorly functional). The function of ALR-K is to convert at higher rates hydroxyphenyl-acetaldehyles produced to hydroxyphenyl-alcohols, such as tyrosol (monophenol route, Fig 1A) and hydroxytyrosol (diphenol route, Fig 1B). However, given the fact that both parental strains, BL21 and HMS174 used in this work, possessed the phenylacetaldehyde dehydrogenase gene *(feaB*), acted both as hydroxyphenylacetic acid producers (Fig 1). This led us to hypothesize that *feaB* must have a faster turn over than the natively existing ADH, which favors under normal condition the conversion of HPAA and DHPAA to the respective acetic acid derivatives [30].

Our data showed that even during high levels of tyrosine production from HMS174-HT^OP^ strain (Fig S5), though in the absence of the overexpressed ALR-K, we saw no increase in HT in the introduced pathway, creating partial pools of the intermediate metabolites, DOPA and tyrosol, with non-equimolar HT produced levels (Fig S5). This led us to further assume that some products were further consumed to an alternative route and not to the HT direction. This course is most likely to involve the function of *feaB* gene product activity, producing the respective acid derivatives and thus removing precursors and substrates that would otherwise lead to HT production. Such acid production may affect the activity of the other enzymes tested, AADC and TYR, as showed by the decrease and or accumulation of their respective products DOPA and tyrosol. Further, involvement of the overexpressed ALR-K resolves the biosynthetic bottleneck problem resulting in equimolar production levels of HT as compared to tyrosine initial levels (Fig 7A). Moreover, when ALR-K was functioning, the levels of DOPA and tyrosol remained minimal showing that the biosynthetic flow from tyrosine to HT was running properly. The negative effect of *feaB* to remove intermediate metabolites from the introduced pathway to *E. coli* through AADC and TYR has been shown before [27, 28, 57], during previous attempts to heterologously produce HT from *E. coli*, which explains our status of data. Suggested deletion of *feaB* gene previously resulted in further increase of HT production.

In contrast to previous attempts to produce HT directly from glucose [27, 28], we utilized different gene sources encompassing different activities and different host strains to show the effect of the coexistence of di- and mono-phenol route. However, by including the overproduction of tyrosine the monophenol route was more advantageous for the HT biosynthesis in *E. coli*. Moreover, the utilization of HMS174 strain as a host over the BL21, presented high capacity for tyrosine production (Fig S5). Nevertheless, although the tyrosine produced in HMS174 was high it did not lead to a proportional increase in HT production. Only when *ALR-K (yahK*) was over-expressed, it led to further increase in HT production indicating that an intermediate which may serve as ALR-K substrate inhibited the HT biosynthesis. The ALR-K enzyme is NAPDH dependent and possesses a broad reductase activity on aldehydes [58] and thus it may decongest any phenylacetaldehyde pooling. ALR-D (yqhD) is also a broad-substrate range aldehyde reductase [59] but did not gave any positive result in our system (data not shown).

The final titer of this metabolic construction was about 22 times higher than that of [27] and 1.3 times higher than that of Chung et al. [28] (Table 3), while it was 12 times less than the precursor tyrosine reported to be produced by the tyrosine overproducer strain [29], a fact that dictates space for further optimization.

**Table 3:**
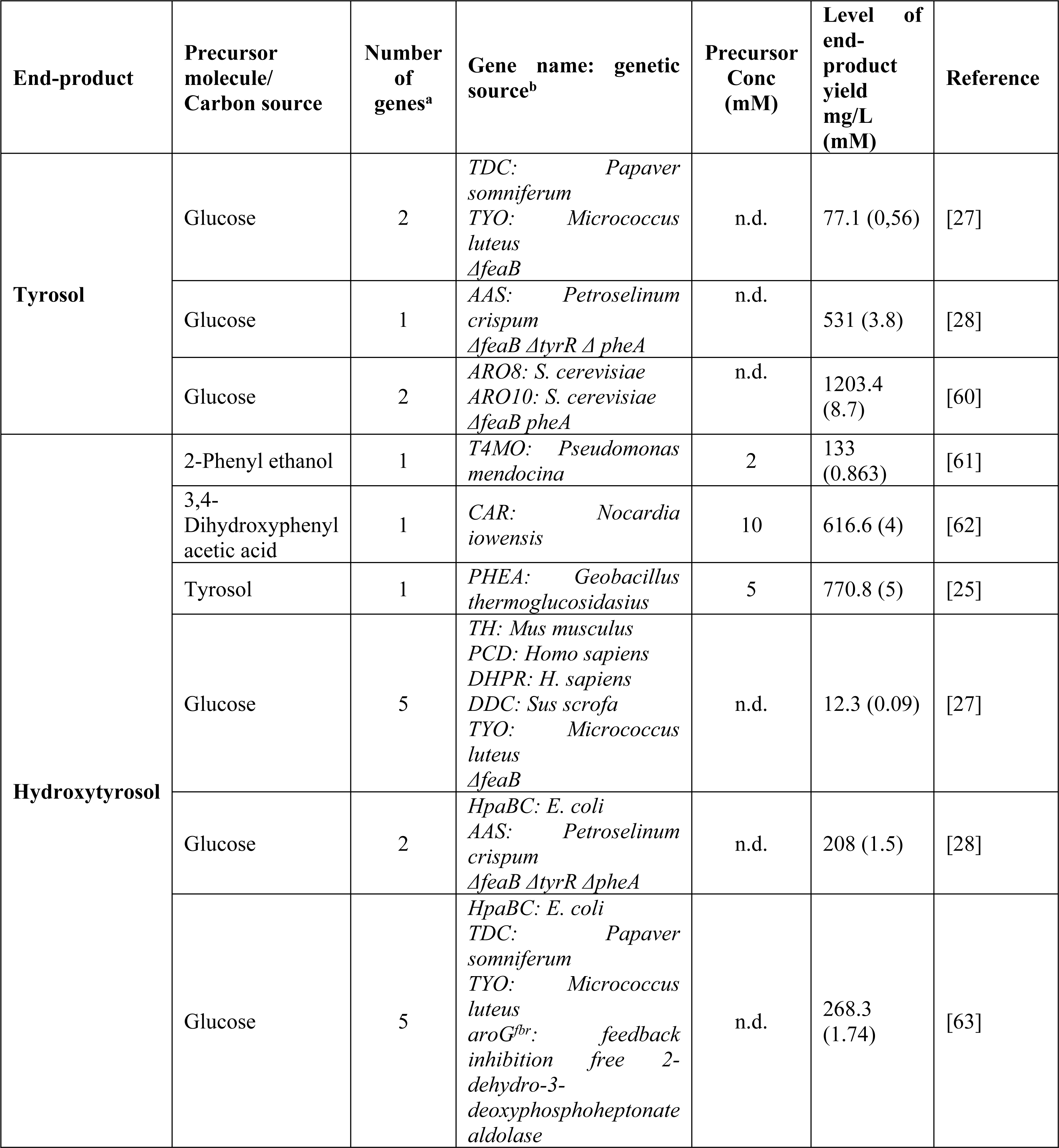
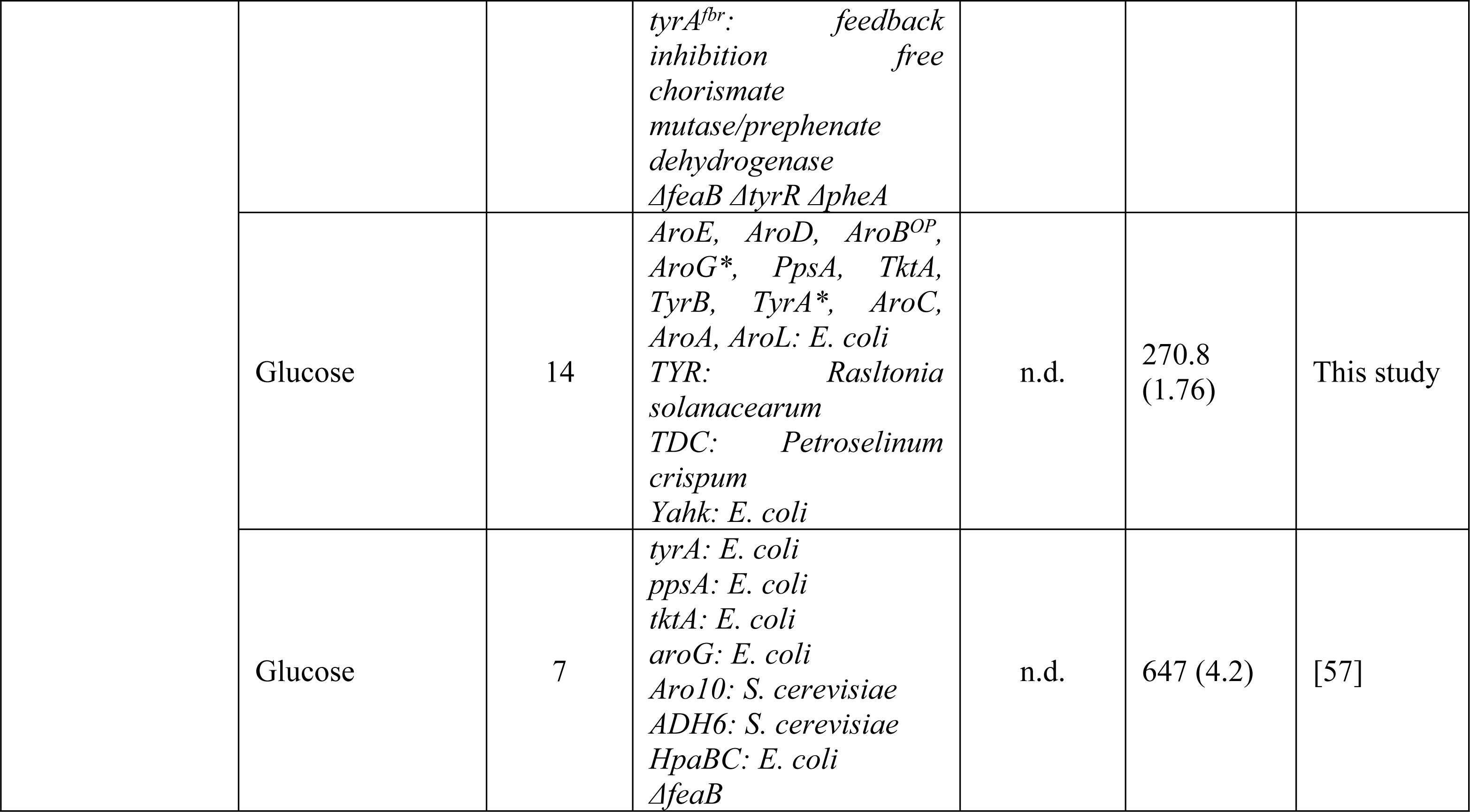
Selected references documenting the microbial production of Tyrosol or Hydroxytyrosol through utilization of microbial biotechnology strategies (n.d., not-determined).

At the time of submission, two new articles drawn our attention for the biosynthesis of hydroxytyrosol in *E. coli*, authored by Li et al. [57] and Choo et al. [63]. The performance of their system is of interest due to the efficacy they achieved, especially for the case of Li et al. [57] who achieved a titer of 647 mg/L (Table 3). They ended up in such a high concentration following a different approach, directing the metabolic flow from 4-hydroxyphenylpyruvate to 4-hydroxyphenylacetaldehyde, tyrosol and finally to HT through the action of a ketoacid decarboxylase, an alcohol dehydrogenase and a 4-hydroxyphenylacetic acid 3-hydroxylase, respectively. The pathway was overloaded with 4-hydroxyphenylpyruvate by the action of an aromatic-amino-acid aminotransferase. Their pathway proved more efficient in terms of HT production, utilizing four enzymes in contrast to our strategy that utilized fourteen; this may be an explanation of why they succeeded higher efficacy.

Concluding, the genetically tractable microbe *E. coli* provides a supreme platform for the combinatorial biosynthesis of plant natural products utilizing heterologous genes from various sources. Microbial production of plant natural products is a promising alternative to traditional methods, sophisticated metabolic engineering of host’s primary and secondary metabolic machinery is needed to improve the final product efficiency or assist in the discovery of novel compounds. Here, we presented the numerous efforts made for the optimization of HT production directly from glucose. Although, the HT concentration reached 270.8 mg/L within 48h, further optimization efforts would be expected to increase the final titers particularly important for the entry of the system to the industrial production.

## Acknowledgements

The authors are grateful to a) Prof. J.D. Keasling (University of California, USA) for kindly providing pS4 and pY3 plasmids, b) Prof. C. B. Voltattorni (Università di Verona) for providing the DDCmut gene variant, and c) emeritus Prof. Nickolas Panopoulos (University of Crete) for providing DNA from the strain GMI1000 of *Ralstonia solanacearum*.

## Authors’ contributions

ET and FV conceived and supervised the project. ET and FV designed the experiments and wrote the manuscript. ET, EN, TP, TN, MH, LS and FV performed the experiments and analyzed the data. All authors read and approved the final manuscript.

## Supporting information

**S1 Fig. Prediction of the presence of a translocation signal peptide on the *Ralstonia solanacearum* Tyrosinase (TYR) N-terminal.**

The PRED-TAT on line software was utilized. The cleavage site was predicted between the two alanines (in bold) of AVAAD (at position 31-32 of the protein sequence).

**S2 Fig. The two plasmid modules named pS4 and pY3 for the reconstruction of L-tyrosine overproducing route from glucose.**

*AroA*, 5-enolpyruvoylshikimate 3-phosphate synthase; *AroB*, Dehydroquinate synthase; *AroC*, chorismate synthase; *AroD*, Dehydroquinate dehydratase; *AroE*, shikimate dehydrogenase; *AroG*, 3- deoxy-D-arabino-heptulosonate synthase; *AroL*, shikimate kinase; *PpsA*, phosphoenolpyruvate synthase; *TktA*, transketolase A; *TyrA*, chorismate mutase/prephenate dehydrogenase; *TyrB*, tyrosine aminotransferase; P, Promoter; O, Operator. T, Termination sequence.

**S3 Fig. Tyrosinase and Aromatic aminoacid decarboxylase overexpression in *Escherichia coli* to certify their expression.**

(A) Upper part, protein expression in *Escherichia coli* to certify the tyrosinase protein expression. In the first lane total proteins from *E. coli* BL21(DE3)-pRSF in non-induced conditions were loaded. In the second, total proteins from BL21(DE3)-pRSF-RsTYR in non-induced conditions were loaded while in the third lane total proteins from BL21(DE3)-pRSF-RsTYR in induced conditions were loaded. The arrow in the protein marker helps to estimate the size of the protein band. Lower part, colorimetric assay with the protein crude extracts to assess the activity of tyrosinase as described in Material and Methods. (B) Protein expression in *Escherichia coli* to certify the AADC expression. In the first lane the PiNK prestained protein ladder was loaded. In the second and the third lanes total proteins from BL21(DE3)-pRSF-PcTDC in induced and non-induced conditions were loaded respectively. The arrows in the protein marker help to estimate the size of the expressed protein band.

